# Region- and layer-specific glutamatergic synapse development in the nascent cortical hierarchy

**DOI:** 10.64898/2026.02.17.706315

**Authors:** Luca Discepolo, James McAllister, Rosie Russell, Sarah Apilado, Gabriella Margetts-Smith, Daniela Franchini, Seth G.N. Grant, Cian O’Donnell, Michael C. Ashby, Paul G. Anastasiades

**Affiliations:** University of Bristol, Translational Health Sciences, Dorothy Hodgkin Building, Whitson Street, Bristol BS1 3NY, UK; Ulster University, Intelligent Systems Research Centre, Derry~Londonderry Campus, BT48 7JL, UK; University of Bristol, School of Psychology & Neuroscience, University Walk, Bristol BS8 1TD, UK; University of Exeter, Exeter EX4 4QH, UK; Genes to Cognition Programme, Institute for Neuroscience and Cardiovascular Research, University of Edinburgh, Edinburgh EH16 4SB, UK; Simons Initiative for the Developing Brain (SIDB), Centre for Discovery Brain Sciences, University of Edinburgh, Edinburgh EH8 9XD, UK

## Abstract

Neocortical synapses are highly dynamic during brain development, undergoing formation, elimination, and maturation before acquiring properties that support adult cognition. Individual neocortical regions develop at different ages and individual layers within these regions contain distinct neuronal subtypes that process unique patterns of local and long-range synaptic input. To better understand the development of the cortical hierarchy we explored the laminar maturation of glutamatergic synapses across cortical regions. Synapse maturation was associated with the upregulation of the postsynaptic density protein PSD95. This maturation occurred in a region- and layer-specific manner — layers associated with feedforward pathways develop earlier, while layers associated with higher-order circuits develop later. Our findings highlight adolescence as an important period for the cortex-wide maturation of synapses in cortical layer 1, synapses known to receive top-down feedback from higher-order cortices. We propose that this delayed adolescent maturation of top-down input represents a global signature of cortical development and seemingly acts as the final stage of outside-in brain maturation.

## Introduction

The anatomical connectivity of the neocortex is organized hierarchically, with primary sensory regions at the bottom and higher-order association cortices at the top (Felleman and Van Essen, 1991; Harris et al., 2019). The canonical view is that the timeline of brain development mirrors this hierarchy, with sensory regions observed to undergo maturation prior to association areas across species (Larsen et al., 2023; Reh et al., 2020). However, there is some evidence from both primates and rodents that challenges this idea, instead suggesting that the timeline of maturation in sensory and association areas is remarkably similar (Bourgeois et al., 1994; Myme et al., 2003). At the mechanistic level, cortical circuit maturation has predominantly been studied in primary sensory areas due to the ease with which sensory afferents can be manipulated (Erzurumlu and Gaspar, 2012; Fox and Wong, 2005). These studies indicate that developmental sequencing follows the canonical circuit (Douglas and Martin, 2004; Harris and Shepherd, 2015), with sensory critical periods occurring first in thalamo-cortical synapses, then between cells within layer (L)4, followed by the ascending L4➔L2/3 pathway, and L2/3 recurrent synapses (Erzurumlu and Gaspar, 2012). This sequential, as opposed to synchronous, maturation follows an “outside-in” rule whereby critical periods occur successively along the arc of sensory information transfer from the periphery to the neocortex (Erzurumlu and Gaspar, 2012; Sehara and Kawasaki, 2011). If, or how, layer-specific maturation occurs in higher-order, associative regions remains to be determined.

Brain development culminates in adolescence when higher-order brain regions, including the prefrontal cortex (PFC), are thought to undergo their final maturation (Anastasiades et al., 2022; Delevich et al., 2019; Larsen and Luna, 2018). This process is key to the acquisition of adult-like cognition (Larsen and Luna, 2018; Pöpplau et al., 2024; Zhu et al., 2024) and likely contributes to the increased susceptibility of the adolescent brain to a range of PFC-associated neuropsychiatric and mental health conditions (Anastasiades et al., 2022; Blakemore, 2019; Solmi et al., 2021). Despite being of clear importance, less is known about when and how PFC maturation occurs compared to sensory areas. Thalamic input to the PFC is present from the first postnatal week in rodents (Ferguson and Gao, 2014). Neural activity during this time is important for proper network formation and function (Bitzenhofer et al., 2021) and depends on preconfigured synaptic parameters (Chini et al., 2024). Subsequent thalamic activity during adolescence orchestrates the emergence of adult-like connectivity and cognition (Benoit et al., 2022; Petersen et al., 2024; Yang et al., 2025). However, it is unclear if the PFC undergoes thalamus-first maturation, mirroring sensory cortices, or if its distinct architecture, with cortical, amygdala, and hippocampal inputs distributed across different layers (Anastasiades and Carter, 2021), gives rise to alternative layer-, or input-specific patterns of development (Kroon et al., 2019; Pattwell et al., 2016).

The laminar separation of neocortical inputs is perhaps most evident in the feedforward and feedback projections that mediate communication between higher-order association and lower-order sensory regions, with feedforward inputs targeting middle layers and feedback connections particularly dense in outer L1 (Felleman and Van Essen, 1991; Harris et al., 2019; Larkum, 2012). Although the thinnest and most cell-sparse cortical layer, L1 displays significant complexity and acts as a key node for the integration of diverse long-range inputs (Huang et al., 2024), which account for about 90% of L1 synapses (Larkum, 2012). These inputs are thought to convey predictions and contextual information, as well as signalling behaviourally relevant outcomes to drive learning and support goal-directed behaviour (Norman et al., 2021; Pardi et al., 2023; Schuman et al., 2021). Despite the importance of both feedback signals and L1 to cortical function, the study of cortical development has largely focused on feedforward pathways (Erzurumlu and Gaspar, 2012; Hooks and Chen, 2020). Consequently, it remains unclear if improvements in cognitive capabilities that depend on feedback connections are driven by the maturation of the frontal areas that are the source of these connections (Nabel et al., 2020), or if sensory areas undergo more protracted development than is typically assumed allowing them to better respond to feedback.

Long-range communication in the neocortex is primarily mediated by glutamatergic synapses, whose number, morphology and molecular composition vary significantly during development (Lohmann and Kessels, 2014), contributing to the formation and maintenance of new memories and the emergence of higher cognitive functions as we age (Fitzgerald et al., 2015; Migaud et al., 1998). Individual synapses are highly dynamic across development (Bhatt et al., 2009; Holtmaat and Svoboda, 2009) but at the population level there is an initial phase of overproduction followed by a period of refinement that removes inactive synapses while stabilizing those that are active (Bourgeois et al., 1994; Grutzendler et al., 2002; Nagappan-Chettiar et al., 2024). At the molecular level, synapse maturation involves transitions in the relative composition of the synaptic proteome, including AMPA and NMDA receptors (Anastasiades and Butt, 2012; Ashby and Isaac, 2011; Frank et al., 2016; Matta et al., 2011; Skene et al., 2017) and scaffold proteins that anchor synaptic signalling machinery at the postsynaptic density (PSD) (Cizeron et al., 2020; Frank et al., 2016; Sheng and Kim, 2011). The membrane-associated guanylate kinase (MAGUK) proteins, which include PSD95 and SAP102, are key components of the PSD (Husi et al., 2000; Sheng and Kim, 2011; Zheng et al., 2011). Individual synapses contain unique combinations of PSD95 and SAP102 (Zhu et al., 2018), which contributes to region and layer-specific synaptic properties and neural dynamics (Hansen et al., 2025). The expression of PSD95 and SAP102 is highly developmentally regulated (Cizeron et al., 2020), yet it is unclear if, or how, this process contributes to the development of specific layers and regions across the cortical hierarchy.

To resolve these questions, this study aims to determine the synaptic maturation of functionally distinct brain regions at different ends of the cortical hierarchy: the primary somatosensory (barrel) cortex (SSp) and the prelimbic subdivision of the medial PFC. Using high-throughput automated detection of fluorescently-tagged MAGUK puncta across cortical layers alongside computational modelling of synapse dynamics, we highlight both similarities and key differences in the laminar maturation profiles of different regions of the neocortex.

## Methods

### Experimental animals

Male PSD95^eGFP/eGFP^; SAP102^mKO2/y^ and female PSD95^eGFP/eGFP^; SAP102^mKO2/mKO2^ (Zhu et al., 2018) or C57BL/6J mice (Charles River, UK) were housed in our animal facility on a 12/12 hour light/dark cycle with *ad libitum* access to food and water. For all experiments the age range for each developmental cohort was ±2 days of the reported age. All animal experiments were performed in accordance with The Animal (Scientific Procedures) Act 1986.

### Histology and immunohistochemistry

Mice received a lethal dose of sodium pentobarbital (200mg/ml) before cardiac perfusion with 0.01M PBS followed by 4% PFA. The brain was then extracted and stored in 4% PFA at 4°C until slicing. Slicing was performed on a Leica VT1000S vibratome. For PSD95 and SAP102 labelling 50 µm sections were taken and mounted on suprafrost slides (Thermo Fisher) with Vectashield mounting media with DAPI (Vector labs) and then coverslipped. For VGlut2 staining slices were cut at 50 µm and underwent antigen retrieval in sodium citrate buffer at 80°C for 30 minutes. Slices were allowed to cool before being washed in 0.01M PBS. Slices were incubated at room temperature for 1 hour in blocking solution: 5% normal goat serum in PBS-T (0.01M PBS and 0.3% Triton-X) with 0.05% sodium azide. Slices were then incubated with guinea pig anti-Vglut2 (1:1000; Synaptic Systems, Cat.No. 135404) in blocking solution overnight at 4°C. Slices were washed in 0.01M PBS before the secondary antibody (Goat anti-Guinea Pig Alexa 647, 1:1000; Invitrogen, A21450) was applied in blocking solution for one hour at room temperature. Slices were washed in 0.01M PBS and mounted on suprafrost slides using ProLong Gold mounting media with DAPI (Invitrogen) and then coverslipped.

### Imaging

All imaging was undertaken at the Wolfson Bioimaging Facility at the University of Bristol. Widefield images were obtained with an Olympus VS200 slide scanning microscope using a 10x (NA 0.4) objective. For region-specific analysis, confocal images were obtained on a Leica SP8 confocal microscope using either a 10x (NA 0.3) or 63x (NA 1.4) objective. For puncta imaging across the depth of the cortex, we captured 5 separate images using the 63x objective. The top of position 1 was aligned to the pial surface with 200 µm spacing between images. For PSD95 and SAP102 imaging, we calibrated microscope settings to allow reliable signal detection across the entire developmental range of the study in advance and applied the same settings across all imaging sessions.

### Puncta and nuclei detection

Puncta and nuclei detection was performed using the Modular Image Analysis (MIA) plugin for ImageJ (Cross et al., 2024). This first involved applying a 2D Gaussian filter (sigma = 2 px) to reduce noise, followed by the sliding paraboloid implementation of background subtraction using a radius of 2 px. Puncta were detected using the Laplacian of Gaussian spot detector in TrackMate (Tinevez et al., 2017) (implemented via MIA) and 2D Gaussian profiles fit to each detected punctum to estimate sigma. Spot intensity was measured for all puncta by taking into consideration all pixels within 2 px of the object.

The same plugin was also used to segment and measure nuclei. This involved applying a 2D median filter (radius = 2 px) followed by application of the sliding paraboloid background subtraction; however, retaining the background image as the ongoing representation of the nuclei. Nuclei were detected using the StarDist plugin (Schmidt et al., 2018) on images scaled in X and Y by a factor of 0.3. These nuclear objects were then converted to a binarized representation, upscaled to the full resolution and subject to a watershed transform to ensure all proximal objects were detected separately (Legland et al., 2016). Any nuclei smaller than 25 μm^2^ or with texture “correlation” metrics less than 0.4 were removed from further analysis (Haralick et al., 1973).

For optimal detection we required parameters that had good sensitivity to detect puncta but also specificity to avoid false positives. To determine detection parameters that reliably identified puncta across all ages of our developmental study we chose a random subsample of images spanning all age groups and manually scored identified puncta across a range of detection settings. We chose settings that gave high detection accuracy across all ages. For each individual punctum the MIA plugin calculated values of mean and max intensity. The 2D Gaussian profiles fit to each detected punctum was taken as a proxy for size.

To control for differences in cell density across development and layers we performed additional analysis of puncta density which controlled for the area taken up by the cell nucleus, which is devoid of PSD95 puncta. For each image we subtracted the total area of the nuclear objects detected using the MIA plugin from the area of the image and used this area value to calculate a corrected puncta density for each image using the formula below.

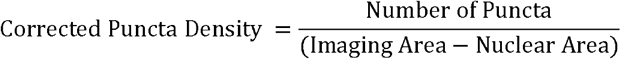

To determine how individual puncta properties contribute to overall PSD95 fluorescence values we took the average 10x fluorescence intensity and compared it to a model based on the calculated puncta number, average intensity and size values on a mouse-by-mouse basis. We used relative maturation values (ranging from 0 to 1) for all calculations, allowing us to determine how a developmental change in the properties of individual puncta parameters contributed to a change in bulk fluorescence. The puncta model was as below:

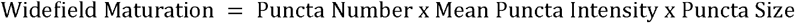

To determine the influence of the different puncta parameters on PSD95 development, either puncta number, intensity, or size was artificially set to zero. The corresponding model error was calculated in each case on a mouse-by-mouse basis and pooled across developmental stage to determine the variable that most strongly impacted model performance.

### Computational modelling

Computational modelling was performed to simulate and analyse the dynamics of synapse formation, maturation, elimination, and synaptic weight. The model considered three possible states of a given synapse: a potential resource pool (*P*), an immature population (*I*), and a mature population (*M*). The numbers of synapses in the resource pool, immature population, and mature population are denoted by *N*_*P*_,*N*_*I*_,and *N*_*M*_, respectively. All synapses start in the potential resource pool but may transition from the resource pool to the immature state (“synapse creation”); from the immature state to the potential resource pool (“synapse elimination”); from the immature state to the mature state (“synapse maturation”); and from the mature state to the immature state (“synapse de-maturation”). These transitions are rate regulated by the creation rate (*c*), elimination rate (*e*), maturation rate (*m*), and de-maturation (*d*).

Population dynamics of synapses was modelled with two different approaches for comparison, using (1) random walks and (2) differential equations.

#### (1) Random Walks

In the random walks version, each synapse is represented by a random variable that can occupy one of the three states *P,I* and *M*. At each discrete time step, a synapse may transition between states based on specific probabilities *p*_*c*_,*p*_*e*_,*p*_*m*_ and *p*_*d*_ (derived from the rates *c,e,m* and *d*) and according to the transitions presented by the transition matrix *T*:

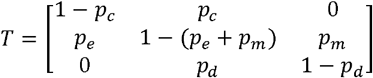

Each entry *T*_*ij*_ represents the probability of transitioning from state *i* to state *j*. For example, the entry *T*_*PI*_ ( = *T*_12_) = *p*_*c*_ denotes the probability of a synapse transitioning from the potential resource pool state to the immature state. This transition matrix summarises the fact that a synapse can transition *P* ⇌ *I* and *I* ⇌ *M* with relevant probabilities, but not directly *P* ⇌ *M*. At every timestep the model calculates the numbers of synapses in each of the three states, giving results for *N*_*P*_,*N*_*I*_, and *N*_*M*_ across time. This Markov chain model is simulated over many trials to calculate the average population dynamics.

The random walks model has the advantage that its stochastic nature captures the randomness inherent in biological processes, providing a relatively realistic simulation of *individual* synapse behaviour. However, it is more computationally expensive and less analytically insightful.

#### (2) Differential Equations

In the differential equations model, the population dynamics are described by the following equations:

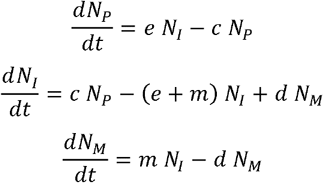

The differential equations model loses the stochastic nature of the individual synapse state transitions, instead modelling *average* population behaviour. However, it is more computationally efficient and lends itself more easily to analysis.

For all synapses in the mature population, we also model synaptic weight dynamics using a Kesten Process, which has been previously found to account for the fluctuations of synapse sizes over time (Statman et al., 2014). This model is a combination of stochastic multiplicative and additive processes, described by the following equation:

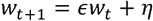

where *W*_*t*_ represents the synaptic weight at time *t*, and *ϵ* and *η* are the multiplicative and additive normal random variables, respectively. At every timestep, a random number is sampled independently for both *ϵ* and *η*, which together cause the synaptic strength to fluctuate stochastically over time. Parameter values were based on Statman et al. 2014. The simulation timestep was set to 0.1 days. The Kesten process with appropriate parameters gives resulting synaptic strength distributions that are broad, skewed and relatively stable over time, yet individual synapse strengths exhibit significant spontaneous fluctuations (Statman et al., 2014). In the differential equations version where *population* dynamics are modelled instead of individual synapses, the model continues in parallel to stochastically simulate individual synapses within the mature population by the Kesten process, ensuring that the number within this simulated *weight dynamics* pool aligns with the population count prescribed by the differential equations.

The model was initially implemented with constant transition rates *c,m,e*, and *d*, with the only added rule that when a synapse transitions from the mature to immature population, the synapse with the smallest synaptic weight is removed from the mature population (this has no impact on the population dynamics). However, mathematical analysis (see appendix) proved that, if all synapses start in the resource pool, then the combined synapse population (*N*_*I*_ *N*_*M*_) increases strictly monotonically with constant values of *c,m,e*, and *d*. To capture an initial increase and then *elimination* of synapses, we proceeded to use variable rates with developmental age.

Variable rates of synapse creation, elimination, and de-maturation are more neurobiologically realistic and give rise to more complex population dynamics. A variable rates model implemented the rates *c* and *e* in a *time*-dependent manner using exponential functions such that the rates start high and decrease to a baseline over developmental time (Fig. S10B). The equations are:

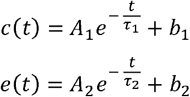

The parameter *b*_*i*_ controls the asymptotic baseline value that the rate decays to; *τ*_i_ is the time constant and controls the timescale of the exponential decay; *A*_*i*_ is the amplitude coefficient, determining the initial magnitude of the decaying component. In this study the rates were set such that *b*_1_ = *b*_2_ = 0.2, ensuring that creation and elimination rates eventually equilibrate, consistent with experimental data (Holtmaat et al., 2005). We also modelled a variable rate of de-maturation, however in a *weight*-dependent manner, for all synapses in the mature population. This can also be modelled simply with an exponential function:

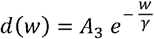

Here *w* is the synaptic weight of a given synapse; *A*_3_ is the amplitude coefficient; and *γ* is the weight decay constant. This part of the model did not include a positive baseline term, to ensure that the probability is vanishing for increasingly large synapses. This captures the observation that biologically, smaller synapses are more likely to de-mature and large synapses are unlikely to de-mature (Holtmaat et al., 2005; Trachtenberg et al., 2002). For the variable de-maturation rate in the differential equations model, the model calculates *d* by taking a weighted expectation through the probability distribution across synaptic weights. Synapses are then removed probabilistically from the simulated *weight dynamics* population (for the Kesten process) to align with the population count *N*_*M*_ computed by the differential equations.

We performed a global sensitivity analysis on the model to quantify how variations in the rate parameters (*A*_1_, *τ*_1_, *A*_2_, *τ*_2_, *m*,, *A*_3_,*γ*) affected the final synapse counts and the timing of peak synapse numbers. We used the regression method in the Julia package `GlobalSensitivity.jl` to perform this analysis, which produced standard regression coefficients for each parameter and outcome of interest.

We calculated the survival fraction of synapses during two developmental stages: early development (PND 16 to 26) and adulthood (PND 70 to 88), corresponding with empirical data in (Holtmaat et al., 2005). To compute the survival fractions, we took the initial synapse count in the combined population at the beginning of each of these developmental stages, and then tracked the states of these synapses, calculating the fraction that survive over the subsequent days relative to the initial count. This was repeated over multiple trials and averaged.

**Table 1.**
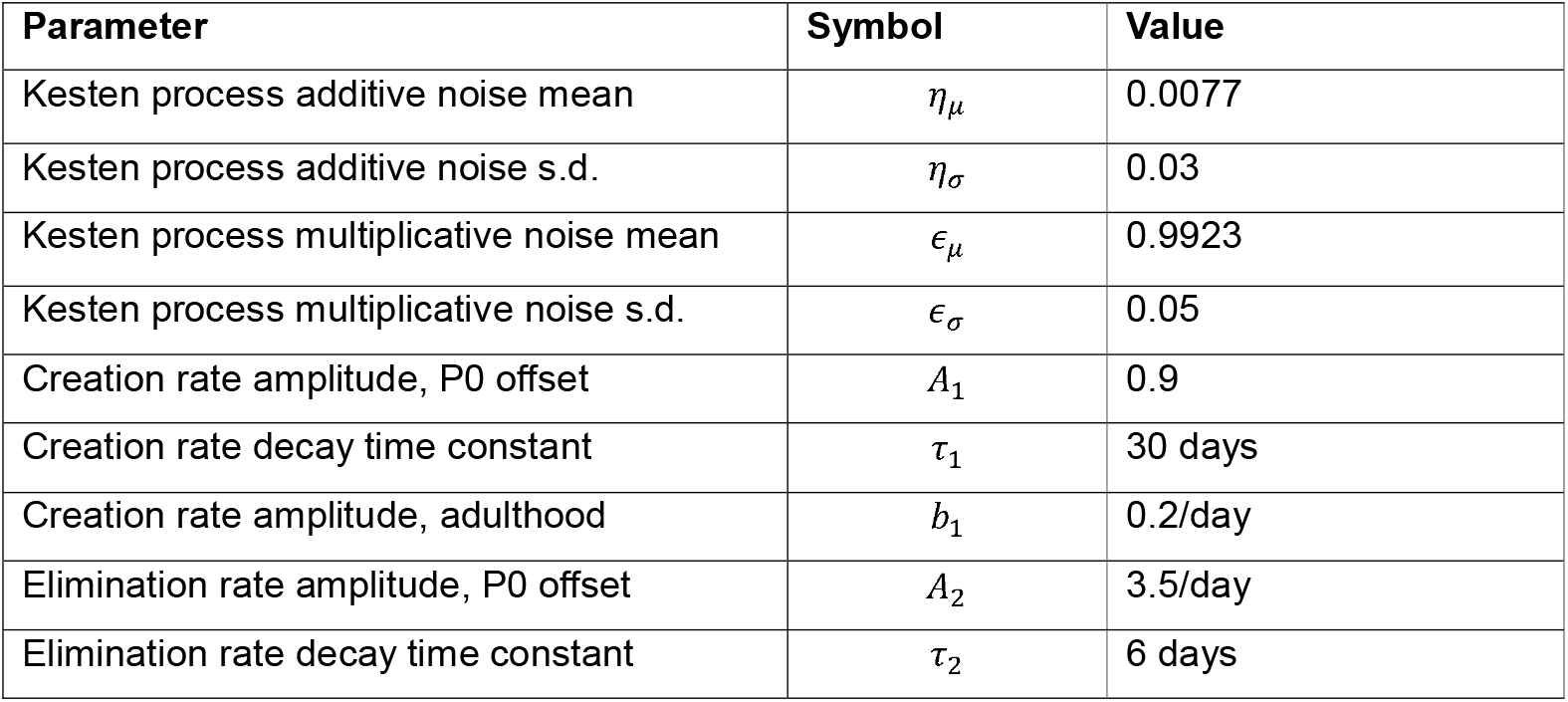
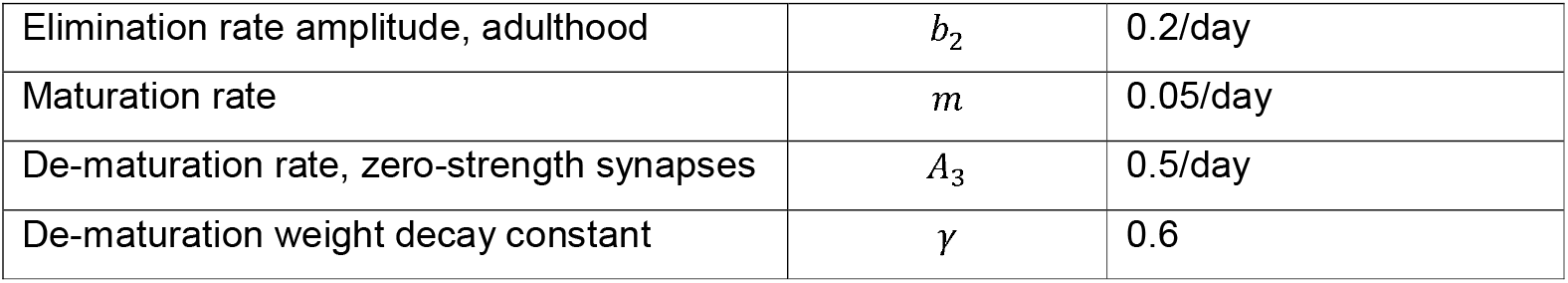
Computational model parameter values.

### Brain atlas alignment

To examine the effect of hierarchical organisation on the adolescent maturation of the cortex, we selected 18 cortical areas spanning the cortical hierarchy. The cortices analysed were the secondary motor (MOs), prelimbic (PL), infralimbic (ILA), medial orbital (ORBm), anterior cingulate area ventral (ACAv), agranular insular area dorsal (AId), primary motor (MOp), primary somatosensory (SSp, focusing on SSpbfd), secondary somatosensory (SSs), ventral retrosplenial (RSPv), dorsal retrosplenial (RSPd), anteriomedial visual (VISam), anterior visual (VISa), dorsal auditory (AUDd), primary auditory (AUDp), posteriormedial visual (VISpm), primary visual (VISp), and anteriolateral visual (VISal). We imaged these areas at the start (P25) and end (P55) of adolescence. Hierarchical location was determined using the corticocortical, corticothalamic, and thalamocortical (CC+CT+TC) hierarchical connectivity scores from the Allen Connectivity Atlas (Harris et al., 2019).

Fluorescent profiles of the 18 cortical areas were obtained in Fiji with the BAR plugin (Ferreira et al., 2015) and background subtracted using off-slice pixel intensity. Profile measurements were recorded from both hemisphere across three slices per brain region. Cortical layers were defined using the ABBA (Aligning Big Brain Atlases) workflow aligning coronal sections to the Allen Mouse Atlas V3p1. Brain alignment was achieved in ABBA (Chiaruttini et al., 2024), uploaded to DeepSlice to register each slice to a comparable atlas section (Carey et al., 2023), with fine adjustments to account for dorsal-ventral and medial-lateral slicing angles performed in QuickNII (Puchades et al., 2019). Annotations were then exported to QuPath (Bankhead et al., 2017) where the registered atlas is overlayed over each image. Layer depths based on the atlas annotations were used to segment fluorescent profiles for relative comparison of average PSD95 and SAP102 fluorescence profiles using custom code in Python.

### Layer assignment in confocal images

For developmental layer assignment from individual images of PFC and SSp we used a combination of VGlut2 staining and DAPI labelling combined with previous studies. VGlut2 staining was performed at P5, P15, P25, P35, P45 and P55. At each age measurements were made across multiple slices and animals to create average layer boundaries in each brain region. Given no changes in cytoarchitecture were observed post P15, we used a single set of layer boundaries for each region thereafter. In PFC, for P15 onwards we used layer assignment based on previous analysis of the distribution of thalamic inputs and projection neurons (Anastasiades et al., 2018), which strongly aligned with our VGlut2 data. At P5 the layer boundaries were determined based on average values of measurements taken across the DAPI and VGlut2 images, with layers scaled from the adult data to account for observed changes in labelling. For SSp, the L1/2 border was determined based on changes in cell density, the start and end of L4 based on the presence of barrels, and the L5/6 border based on the location of the second peak of VGlut2 labelling in deep layers. From P15 onwards we used a single set of layer boundaries in SSp based on the grand average of the P15 to P55 data.

### Data analysis and statistics

Data are reported as mean averages +/- SEM, unless otherwise stated. Comparisons between paired data in individual brain regions or mice was performed using a paired t-test. Developmental comparisons were performed using 2-way ANOVA with Tukeys multiple comparison correction. Comparisons between multiple groups were performed with 1-way ANOVA or Friedman test with correction for multiple comparisons. K-means clustering was performed in Python 3. Spline fitting was performed using a smoothing spline in Prism 10. Statistical analysis was performed in Prism 10 or Python 3. Computational modelling and simulations were performed using the Julia programming language (version 1.11.1).

## Results

### Postnatal development of MAGUK proteins across cortical regions

To study the synaptic maturation of the neocortex in a region- and layer-specific manner, we employed a transgenic mouse line where endogenous MAGUK proteins PSD95 and SAP102 are genetically tagged with the fluorescent proteins eGFP and mKO2, respectively (Zhu et al., 2018). We focused our analysis on SSp and the PFC, allowing us to directly compare the maturation of a primary sensory area and a higher-order cognitive area (Fig. 1A-C). We imaged each area in animals aged postnatal day (P)5 until 4 months (P120), a timespan encompassing multiple developmental stages (Fig. 1D). To compare how the expression of PSD95 and SAP102 changes across development, we imaged brain slices and calculated the average fluorescence across the depth of cortex. We subsequently normalized each value by those observed in the adult (P90) to give a relative maturation score, such that 1 represents the average adult expression level and 0 represents a failure to detect meaningful levels of PSD95 or SAP102. This normalization allowed us to more easily compare the timeline of development for the MAGUK proteins, within and between areas. At P5, the relative maturation of SAP102 was significantly higher than PSD95 across both regions (Fig. S1). SAP102 levels continued to increase, peaking during adolescence before stabilising in adulthood (Fig. 1E-G). By contrast, PSD95 expression increased steadily throughout development, reaching stable values by P55 (Fig. 1H-J), towards the end of adolescence. The ratio of SAP102/PSD95 expression reduced considerably in both regions as development progressed (Fig. S1), consistent with developmental shifts in SAP102 and PSD95 levels in the maturing brain (Zheng et al., 2011).

**Figure 1.**
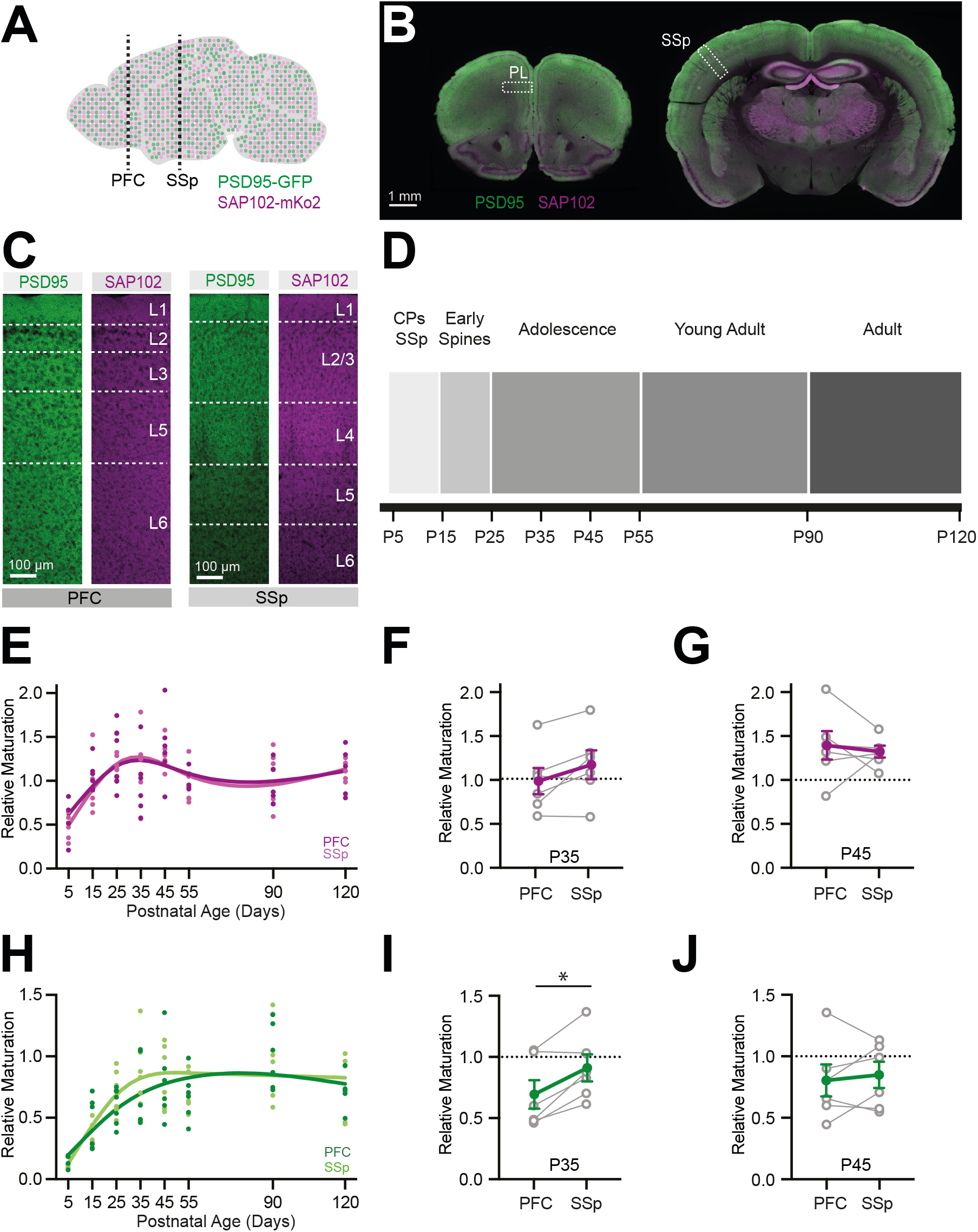
A) Schematic showing transgenic mouse line and regions of interest. B) Example coronal sections showing the regions of analysis in the prelimbic (PL) PFC (*left*) and somatosensory barrel cortex (SSp) (*right*). C) Example images showing the distribution of PSD95 and SAP102 labelling across the laminar depth of the PFC (*left*) and SSp (*right*). D) Developmental timeline showing key periods in the maturation of the neocortex. CPs = critical periods. E) Relative maturation of average SAP102 expression in PFC and SSp normalized to adult levels (P90). Points represent individual mice with fit line to the individual data points. F) Relative SAP102 maturation in PFC and SSp at postnatal day (P)35 showing pairwise, within-mouse comparisons. G) Relative SAP102 maturation in PFC and SSp at postnatal day (P)45 showing pairwise, within-mouse comparisons. H-J) As **E-G** but for PSD95. Number of mice per age group: PFC, P5 = 5, P15 = 6, P25 = 7, P35 = 6, P45 = 6, P55 = 6, P90 = 6, P120 = 5; SSp, P5 = 5, P15 = 6, P25 = 7, P35 = 6, P45 = 6, P55 = 6, P90 = 6, P120 = 5. Values are mean average ± SEM, with exception of data in E & H which show smoothing spline fit to the individual data points. * = p < 0.05.

Although there was some variability in the fluorescence values between mice at any given age (see also (Cizeron et al., 2020)), there was strong intra-mouse correlation in regional fluorescence levels (Fig. S1). To establish if the two areas matured at different rates, we compared the relative fluorescence in SSp and PFC within individual mice. For SAP102, the relative maturation during adolescence was similar across both brain areas at both P35 (SAP102 P35: PFC 0.98 ± 0.1, SSp 1.16 ± 0.2; p = 0.08) and P45 (SAP102 P45: PFC 1.38 ± 0.2, SSp 1.31 ± 0.1; p = 0.6). This was not the case for PSD95, with the relative maturation during mid-adolescence (P35) significantly higher in SSp than PFC, indicating an earlier developmental timeline in somatosensory vs prefrontal areas (Fig. 1I) (PSD95 P35: PFC 0.69 ± 0.1, SSp 0.90 ± 0.1; p = 0.01). This difference had disappeared at P45 during late adolescence (Fig. 1J) (PSD95 P45: PFC 0.8 ± 0.1, SSp 0.84 ± 0.1; p = 0.6). These data indicate that PSD95 maturation is delayed in PFC compared to SSp.

### Defining cortical layers across development

To study layer-specific maturation, it was first necessary to determine how the cytoarchitecture of each cortical region changes during the protracted developmental period of our study. We stained for the presynaptic vesicular glutamate transporter VGlut2, which labels thalamic terminals in the neocortex (Hur and Zaborszky, 2005). Throughout development, our staining mirrored known thalamic innervation patterns in each region (Fig. S2). To determine changes in cortical thickness and laminar cytoarchitecture between P5 and P55, we calculated the distance from the pial surface to either the peak of the L3 VGlut2 band, which corresponds to the peak of mediodorsal thalamus innervation in PFC (Collins et al., 2018), or the start of the L4 VGlut2 band, which delineates the top of barrels in SSp (Sehara et al., 2010)(Fig. S2). There was a significant increase in these values between P5 and P15 in both regions, but little change thereafter (Fig. S2). In addition, we performed automated counts of DAPI labelled nuclei across the depth of PFC and SSp. Consistent with the cortical expansion observed in our VGlut2 data, and known periods of developmental apoptosis in the neocortex (Wong and Marín, 2019), we observed a significant reduction in cell density at all cortical depths between P5 and P15, after which values stabilized (Fig. S3). Using these data and previously published values for cortical layers in the adult (see methods), we were able to assign PSD95 and SAP102 labelling to individual layers across development. Our ability to define thalamo-recipient layers across development allowed us to assess if PFC and SSp regions develop in a thalamus first manner, or not.

### Distinct layer- and region-specific development in SSp and PFC

We first focused our analysis on SSp, whose development is layer-specific and whose sensory critical periods terminate around P20 (Erzurumlu and Gaspar, 2012; Lendvai et al., 2000; Stern et al., 2001). At the level of raw fluorescence, both PSD95 and SAP102 expression showed a significant developmental increase across the different layers of SSp (Fig. 2A,B) (2-Way ANOVA. PSD95 layer p = <0.0001, age p = <0.0001; SAP102 layer p = <0.0001, age p = <0.0001). To analyse the relative trajectory of expression in each layer, we took the individual fluorescence traces, which represent the distribution of PSD95 and SAP102 across cortical layers, and peak normalized them to the largest intensity value in each trace. This normalization allowed us to directly compare the laminar distribution of PSD95 and SAP102 at different ages (Fig. 2C,D) while accounting for changes in the total PSD95 expression across development (Fig. 2A,B). To control for differences in layer thickness, both between layers and across development, we also calculated the normalized labelling density, which takes the average normalized intensity values and divides them by the thickness of each layer at each age. This allows us to assess how enriched PSD95 and SAP102 are in individual layers at different ages. At P5 we observed enrichment of PSD95 and SAP102 staining in granular and infragranular layers that overlap with observed VGlut2 staining patterns (Fig. S2), suggesting early synapse maturation in layers associated with nascent thalamic innervation (Daw et al., 2007; Marques-Smith et al., 2016). For PSD95, we observed that this initial bias towards thalamo-recipient layers decreased over development, with a significant upregulation of relative PSD95 labelling in supragranular layers throughout adolescence (Fig. 2C), such that by adulthood the greatest density of PSD95 signal was in L1 and L2/3 (2-Way ANOVA. PSD95 layer p = <0.0001, age p = 0.028). SAP102 levels were enriched within L4 of SSp at all ages (Fig. 2D) with no developmental shift in their distribution (2-Way ANOVA. SAP102 layer p = <0.0001, age p = 0.79), in marked contrast to the adolescent upregulation of PSD95 observed in superficial layers. These findings are consistent with thalamus first development and also highlight how feedforward (thalamic) and feedback (L1) synapses mature at different rates in SSp.

**Figure 2.**
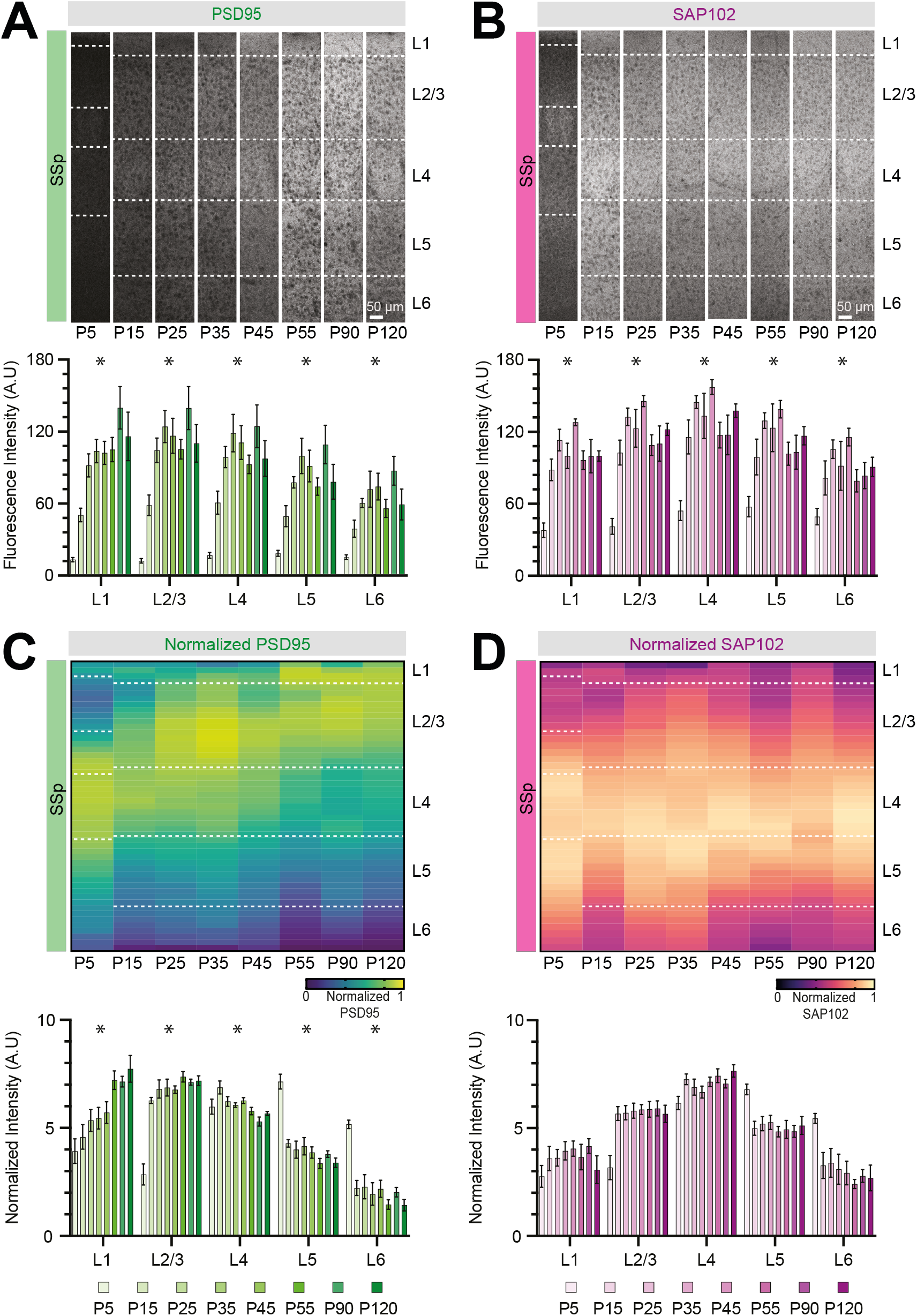
A) (*Top*) Representative images showing the distribution of PSD95 across SSp layers at different postnatal ages. (*Bottom*) Quantification of raw PSD95 fluorescence values for each layer of SSp across development. B) (*Top*) Representative images showing the distribution of SAP102 across SSp layers at different postnatal ages. (*Bottom*) Quantification of raw SAP102 fluorescence values for each layer of SSp across development. C) (*Top*) Heat maps showing the average peak normalized distributions of PSD95 across SSp layers at different postnatal ages. (*Bottom*) Quantification of normalized PSD95 fluorescence density for each layer of SSp across development. D) (*Top*) Heat maps showing the average peak normalized distributions of SAP102 across SSp layers at different postnatal ages. (*Bottom*) Quantification of normalized SAP102 fluorescence density for each layer of SSp across development. Number of mice per age group: SSp, P5 = 5, P15 = 6, P25 = 7, P35 = 6, P45 = 6, P55 = 6, P90 = 6, P120 = 5. All values are shown mean ± SEM. * = layers showing a significant change across development (p < 0.05).

In PFC, PSD95 and SAP102 expression also showed a significant developmental increase across individual layers (Fig. 3A,B) (2-Way ANOVA. PSD95 layer p = <0.0001, age p = <0.0001; SAP102 layer p = <0.0001, age p = 0.001). We again peak normalized the synaptic protein distributions and calculated the normalized labelling density to directly compare their laminar profiles across development (Fig. 3C,D). As in SSp, PSD95 labelling density varied by layer, showing enrichment in thalamo-recipient L1 and L3 and lower levels in L6 (Fig. 3C,D). Surprisingly, the distribution of PSD95 intensity was remarkably consistent between layers, with no significant effect of age (2-Way ANOVA. PSD95 layer p = <0.0001, age p = 0.28). SAP102 labelling showed a distinct laminar profile, being greatest in L2-5 and least in L1 and L6 with again no change across development (2-Way ANOVA. SAP102 layer p = <0.0001, age p = 0.75). These data suggest that although PFC undergoes significant glutamatergic synapse maturation during the first months of postnatal life, this occurs in a layer-independent, rather than a “thalamus first” manner.

**Figure 3.**
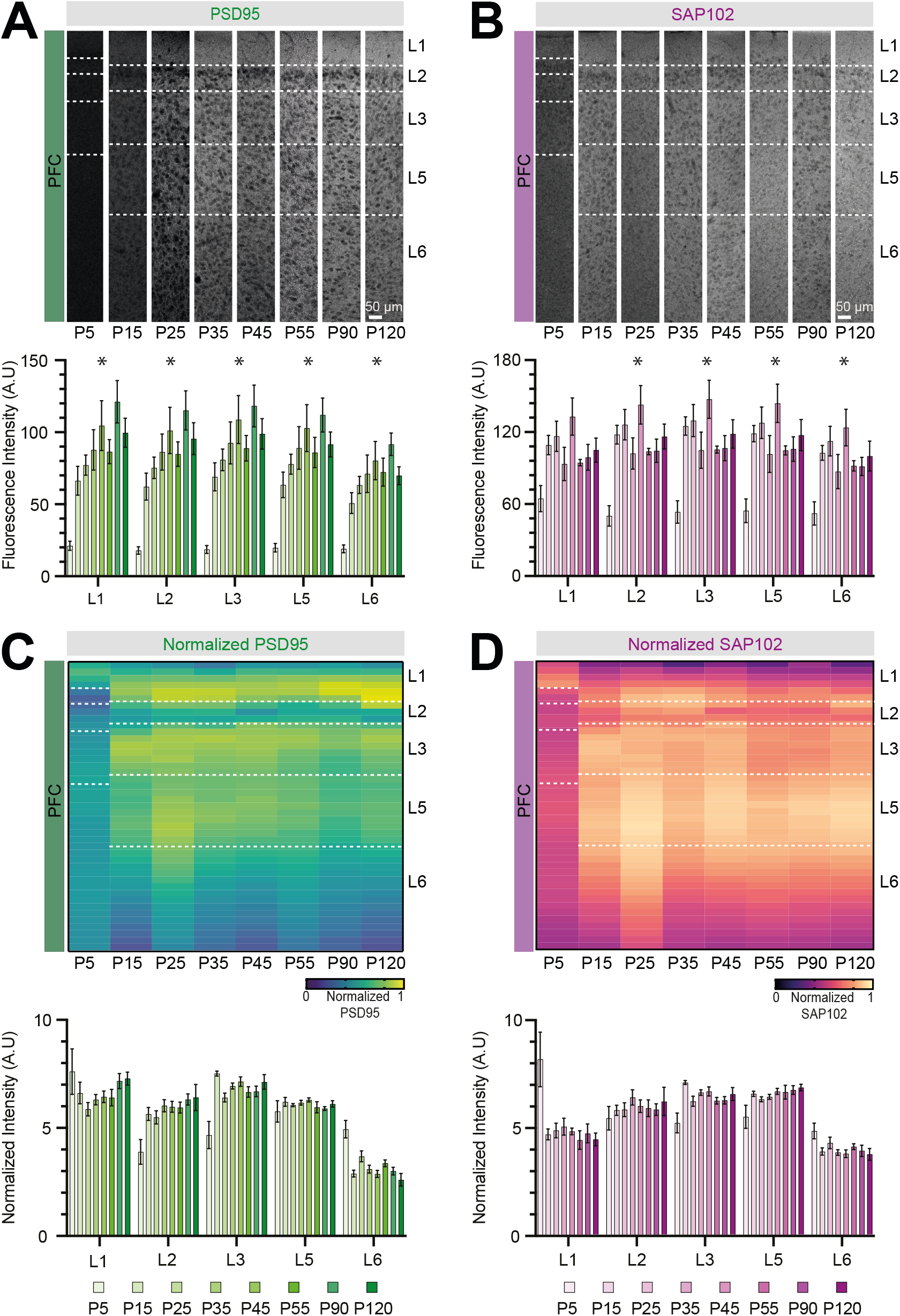
A) (*Top*) Representative images showing the distribution of PSD95 across PFC layers at different postnatal ages. (*Bottom*) Quantification of raw PSD95 fluorescence values for each layer of PFC across development. B) (*Top*) Representative images showing the distribution of SAP102 across PFC layers at different postnatal ages. (*Bottom*) Quantification of raw SAP102 fluorescence values for each layer of PFC across development. C) (*Top*) Heat maps showing the average peak normalized distributions of PSD95 across PFC layers at different postnatal ages. (*Bottom*) Quantification of normalized PSD95 fluorescence density for each layer of PFC across development. D) (*Top*) Heat maps showing the average peak normalized distributions of SAP102 across PFC layers at different postnatal ages. (*Bottom*) Quantification of normalized SAP102 fluorescence density for each layer of PFC across development. Number of mice per age group: PFC, P5 = 5, P15 = 6, P25 = 7, P35 = 6, P45 = 6, P55 = 6, P90 = 6, P120 = 5. All values are mean ± SEM. * = layers showing a significant change across development (p < 0.05).

### Adolescent changes in PSD95 levels vary across the cortical hierarchy

The PFC and SSp both showed changes in MAGUK expression across adolescence, but there were clear region-specific differences in their laminar maturation. To extend these findings, we next sought to determine how other cortical regions across the hierarchy develop. To do so we performed whole-brain imaging of PSD95 and SAP102 pre (P25) and post (P55) adolescence (Fig. 4A). Our hypothesis was that layer-invariant maturation would feature in cortical regions towards the top of the hierarchy, while the delayed maturation of supragranular layers observed in SSp would predominate in lower-order cortices. At each age we aligned images to the Allen Brain Atlas analysing 18 regions spanning the cortical hierarchy (Fig. 4A,B) (Harris et al., 2019). By comparing the change in fluorescence intensity between P25 and P55 we were able to determine how the laminar maturation varied in regions with different hierarchical positions and distinct functional roles within the cortical network. We focused our analysis on PSD95 as we observed minimal adolescent change in SAP102 levels (Fig. S4A). Calculating the absolute change in PSD95 between P25 (n=9) and P55 (n=11) revealed a broad, cortex-wide increase with age (Fig. 4C), which was largest in superficial layers (Fig. S4). The absolute change in PSD95 levels was correlated with the cortical hierarchy in L1 (L1: slope = −4521, p = 0.04), but not other layers (L2/3: slope = −2143, p = 0.2; L4: slope = 2312, p = 0.2; L5: slope = 1280, p = 0.3; L6: slope = 397, p = 0.7) (Fig. 4D), consistent with the adolescent increase we observed in superficial layers of SSp.

**Figure 4.**
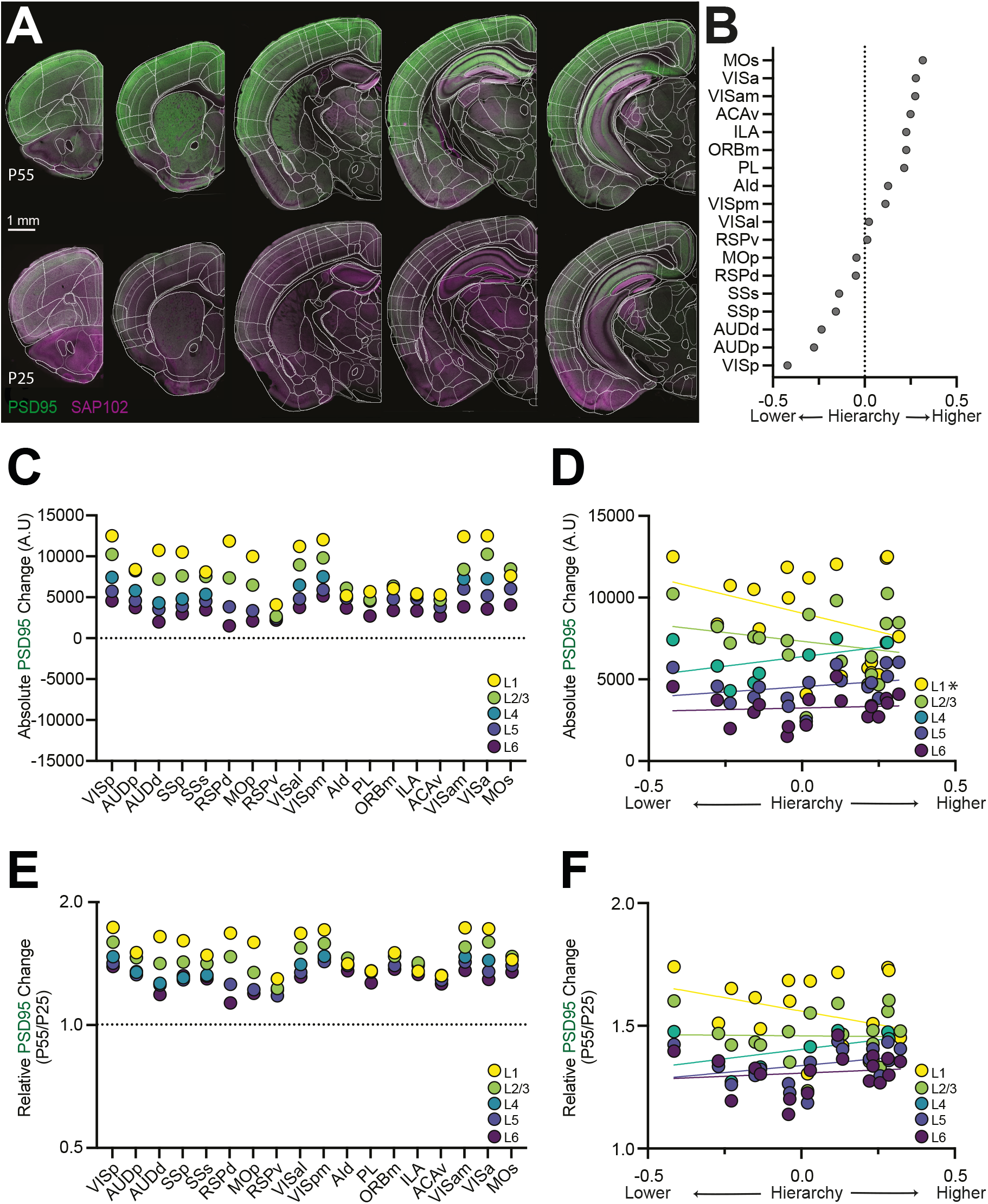
A) Representative images showing PSD95 (green) and SAP102 (magenta) labelling across different mouse brain slices spanning the cortical hierarchy at P55 (top) and P25 (bottom). Overlay shows the Allen Brain Atlas. B) Hierarchy score for the different analysed cortical regions based on cortical and thalamic connectivity (CC+TC+CT) from (Harris et al., 2019). C) Absolute adolescent change in PSD95 fluorescence levels calculated by subtracting the P55 values by the P25 values. Data points show average change for individual layers across the cortical regions shown in **B**. D) Correlation of the absolute adolescent change in PSD95 expression with the hierarchy position of each cortical area. Data is shown for each cortical layer for each of the regions shown in **B**. E) Relative adolescent change in PSD95 fluorescence levels calculated by dividing the P55 values by the P25 values. Data points show average change for individual layers across the cortical regions shown in **B**. D) Correlation of the relative adolescent change in PSD95 expression with the hierarchy position of each cortical area. Data is shown for each cortical layer for each of the regions shown in **B** Number of mice per group: P25 = 9, P55 = 11. Values are shown as mean average. SEM are omitted for clarity. * = p < 0.05.

The absolute change in PSD95 levels indicates how much individual layers change but is not ideal for comparing between structures with different baseline levels of PSD95 expression at P25. To resolve this, we calculated two additional measures. If our hypothesis was correct, in lower-order areas superficial layers should mature more between P25 and P55 than deep layers, while in higher-order areas the layers should mature to the same extent. To test this, we calculated the ratio of PSD95 intensity between L6 and other layers, which also controls for inter-slice labelling variability, and compared this ratio at the two ages (Fig. S4). Importantly, this new dataset confirmed our previous results, with PSD95 expression in the prelimbic PFC maturing in a layer-independent manner, while SSp showed greater supragranular maturation during adolescence (Fig. S4C). When correlating the change in L6 ratio against the cortical hierarchy, there was a significant effect for superficial layers L1 (L1: slope = −0.27, p = 0.001) and L2/3 (L2/3: slope = −0.11, p = 0.04) but not layer 5 (L5: slope = 0.02, p = 0.6). However, when we performed similar analysis for the relative change in PSD95 across adolescence there was no significant effect across any layer (L1: slope = −0.22, p = 0.09; L2/3: slope = −0.01, p = 0.9; L4: slope = 0.15, p = 0.17; L5: slope = 0.11, p = 0.18; L6: slope = 0.05, p = 0.56) (Fig. 4E,F).

Although these data suggest that changes in the cortical circuit may be linked to the hierarchy, we observed significant variability at different hierarchical positions. For example, RSPv, which sits towards the middle of the hierarchy, shows little layer-specificity while higher-visual areas such as VISa and VISam mature in a layer-specific manner (Fig. 4C,D & S4). This led us to ask if the distribution of PSD95 across individual layers, which also varied considerably between brain region (Fig. S5), may inform how individual brain regions change during adolescence. To explore this, we performed unbiased cluster analysis of the raw PSD95 distributions. This revealed two clusters, with cluster 1 dominated by 12 sensory-motor regions, including primary and higher sensory areas, while cluster 2 contained 6 higher-order association cortices (Fig. 5A,B). The segregation of brain regions into these two clusters was similar at P25 and P55 suggesting they represent a fairly static feature of the connectome (Fig. 5A & S5). Comparing the relative adolescent change in PSD95 levels between the clusters revealed that cluster 1 displayed a greater increase in L1 (PSD95 P55/P25 ratio: cluster 1 = 1.62 ± 0.03, cluster 2 = 1.37 ± 0.03; p = <0.0001) and L2/3 (PSD95 P55/P25 ratio: cluster 1 = 1.49 ± 0.02, cluster 2 = 1.37 ± 0.04; p = 0.018) compared to association regions in cluster 2 (Fig. 5C), an effect not observed when comparing deep layers (Fig. S5). This suggests that function, rather than hierarchy may be the main driver of the observed differences across regions. The hierarchical effects we observe may instead be explained by the enrichment of association cortices towards the top of the hierarchy. To confirm this, we performed an analysis restricted to visual areas that span the hierarchy (Fig. 4B) (Harris et al., 2019). This revealed no correlation in any layer (L1: slope = −31, p = 0.99; L2/3: slope = −1159, p = 0.63; L4: slope = −148, p = 0.98; L5: slope = −55, p = 0.98; L6: slope = −983, p = 0.54) (Fig. 5D). These findings suggest that cortical laminar development is strongly linked to function, rather than simply position in the cortical hierarchy. Interestingly, differences in synaptic properties in the two clusters are already present at P25 (Fig. 5A & S5) and are further strengthened via the process of superficial layer synapse maturation during adolescence (Fig. 5B & S5) suggesting that enrichment of L1 synapses in sensory motor regions is a core feature of their functional connectome that matures during adolescence.

**Figure 5.**
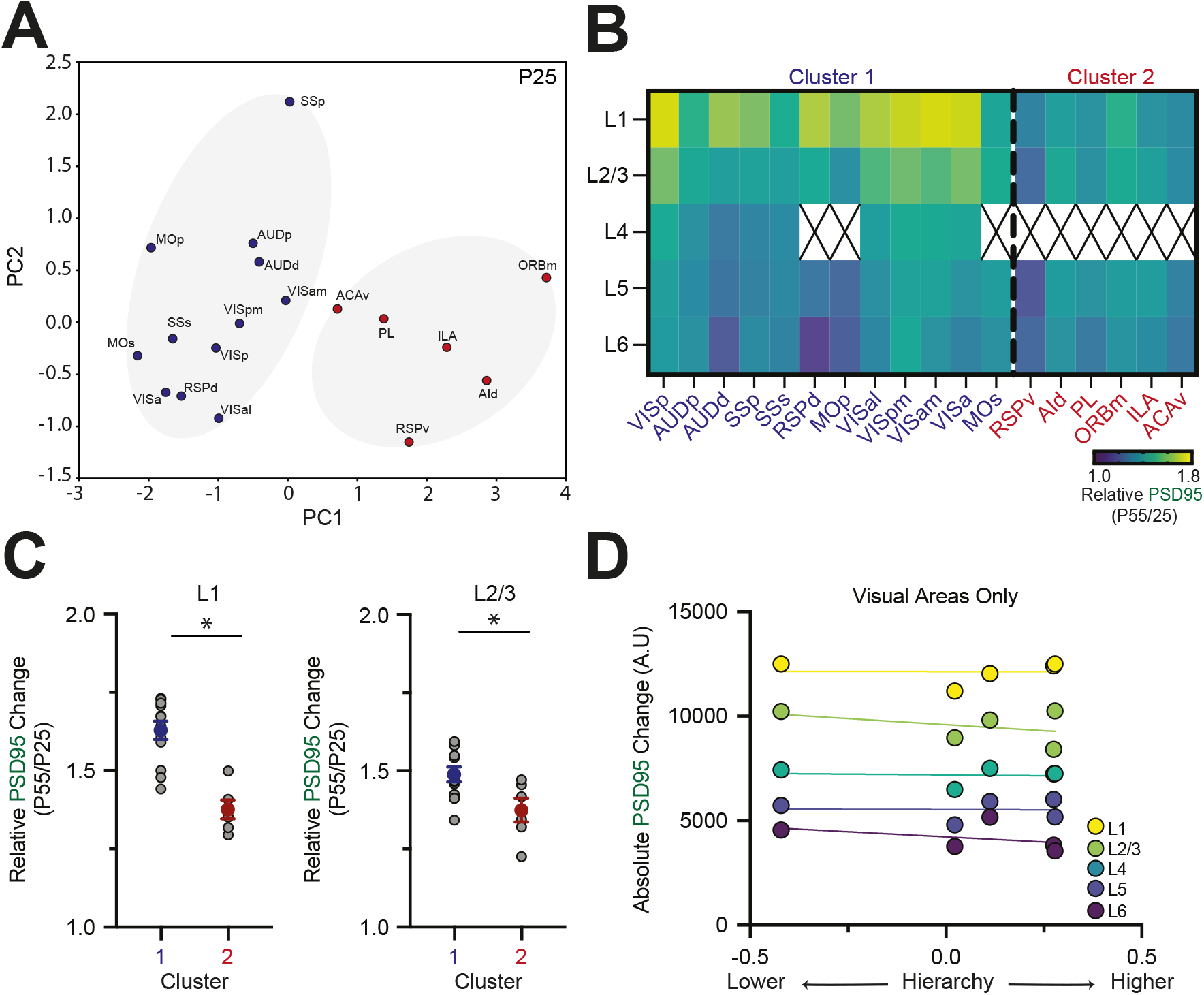
A) Principal component plot highlighting two clusters based on raw PSD95 expression levels at P25. Cluster 1 (blue) contains primarily sensory-motor regions and cluster 2 (red) contains mostly higher-order association cortices. B) Heatmap showing the relative adolescent PSD95 change by layer and regions sorted based on the two clusters in **A**. X indicates agranular cortex lacking L4. C) Comparison of the relative adolescent change in L1 (*left*) and L2/3 (*right*) PSD95 signal between cluster 1 and cluster 2. D) Correlation of the relative adolescent change in PSD95 expression with the hierarchy position of individual visual cortical areas. Number of brain regions per group: Cluster 1 = 12, Cluster 2 = 6. Values are shown as mean + SEM. SEM are omitted in B & D for clarity. * = p < 0.05.

### Postnatal development of PSD95 synaptic puncta

Our analysis up until this point has been performed on bulk fluorescence levels and therefore does not inform about the synaptic mechanisms that drive these changes. Bulk fluorescence signals are a product of the number, size, and fluorescent labelling intensity of individual synaptic puncta. Therefore, to determine how specific synaptic features contribute to the observed developmental changes in PSD95 levels, we performed high-magnification confocal imaging across the cortical depth and applied an automated detection algorithm to segregate individual puncta (Fig. S6). For initial analysis we pooled data across layers and normalized to the adult values, allowing us to compare with bulk fluorescence data (Fig. 1). At P5 we observed a very small number of puncta in both regions (Fig. 6A,B). To confirm that this was not due to insufficient excitation of the fluorophore, we also imaged P5 slices at 2x and 2.5x laser power. Although this yielded a slight increase in puncta number, it was still well below levels observed at P15 with the 1x laser power, ruling out a significant “hidden” pool of puncta at this time (Fig. S6D). From P15 onwards we observed pronounced increase in synapse number in both PFC and SSp, with a similar, yet smaller shift in puncta size and mean intensity (Fig. 6B-D). To determine which puncta parameters best contributed to the development of bulk fluorescence, we modelled the change in PSD95 fluorescence as a product of puncta density, mean intensity and size (see methods). This approach closely approximated the developmental profile of the bulk fluorescence data in both PFC and SSp, as expected (Fig. S6E). Comparing the full model to a reduced model where one of the variables was removed revealed that the main contributors to the development of PSD95 signal were changes in puncta number and intensity, with size having a relatively small effect (Fig. 6E) (median PFC error: number 0.22, size 0.03, intensity 0.17; number vs size p = <0.0001, intensity vs size p = <0.0001, number vs intensity p = 0.86; median SSp error: number 0.17; size 0.03, intensity 0.15; number vs size p = <0.0001, intensity vs size p = <0.0001, number vs intensity p = 0.99).

**Figure 6.**
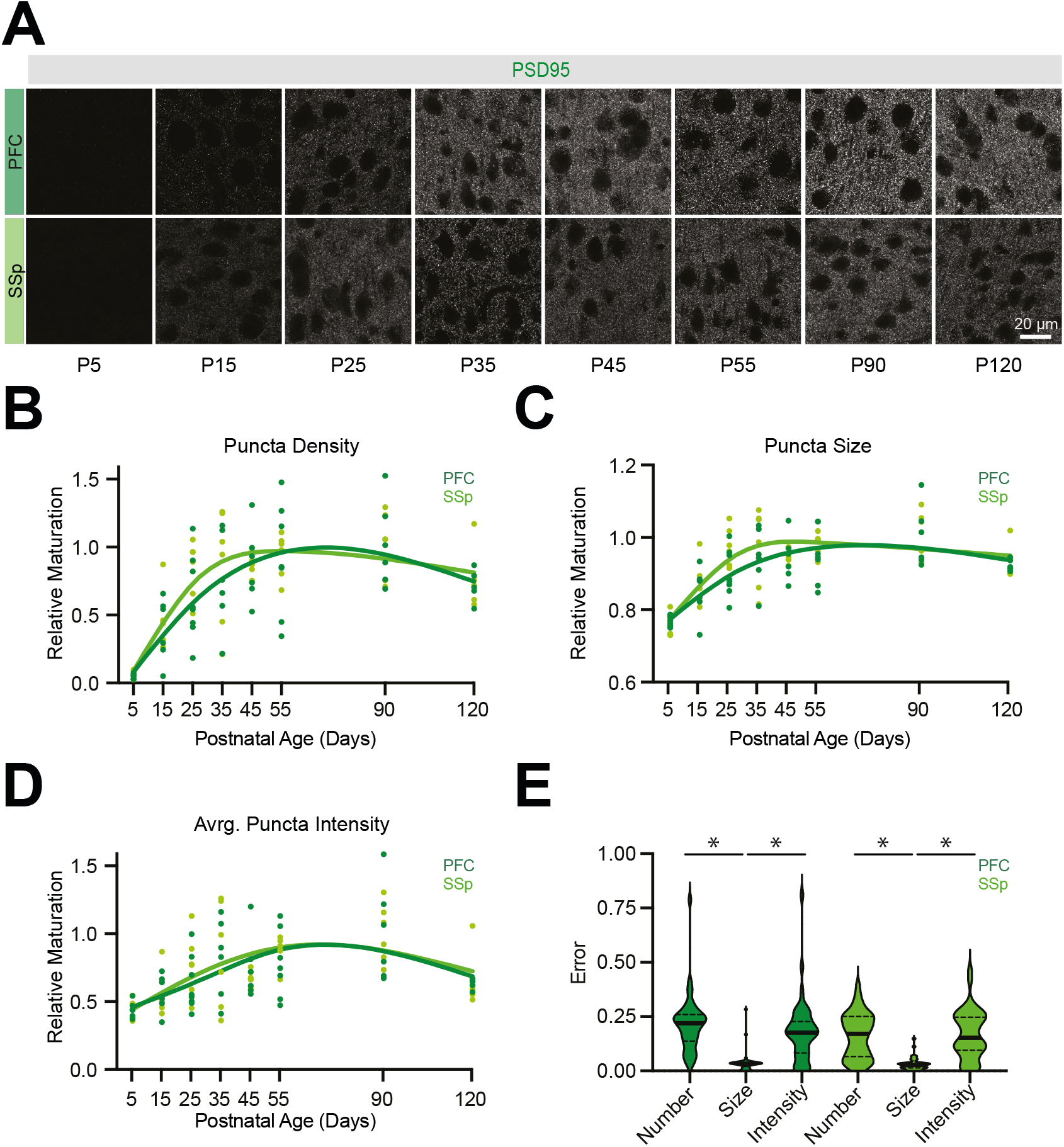
A) Representative high magnification images showing example PSD95 puncta in PFC (*top*) and SSp (*bottom*) at different developmental ages. B) Relative maturation of average puncta density in PFC and SSp normalized to adult levels (P90). Points represent the average puncta density across the cortical depth for individual mice with fit to the individual data points. C) As **B** but for average puncta size. D) As **B** but for average puncta intensity. E) Absolute model error values, representing the change in predicted relative maturation after subtracting either number, size, or intensity data from the model of PFC or SSp development in **Fig. S6E**. Number of mice per age group: PFC, P5 = 6, P15 = 6, P25 = 8, P35 = 6, P45 = 6, P55 = 7, P90 = 6, P120 = 5; SSp, P5 = 6, P15 = 6, P25 = 6, P35 = 6, P45 = 6, P55 = 6, P90 = 6, P120 = 5. Values are shown as mean average with smoothing spline fit to the individual data points (B-D), or median with quartiles (E). * = p < 0.05.

To assess laminar changes in PSD95 puncta, we imaged across the cortical depth using 5 equidistant positions (Fig. 7A,D), with position 1 closest to the pial surface and position 5 the white matter (see Fig. S7 for layer alignment). In the adult PFC, the highest synapse density was observed in superficial layers (Fig. 7A). Because individual layers have differences in cell density (Fig. S3) and synapses are absent from the cell soma (Fig. 5) we performed additional puncta density analysis to account for differences in cell density (see methods). In these data the lack of nuclei in L1 and high density in L2/3 cause a slight shift in the PFC values (Fig. S8), with the biggest increase in puncta density in position 2, which overlaps with ascending thalamo-cortical input from the mediodorsal thalamus (Collins et al., 2018). The relative development of PFC was found to be layer-independent, with the maturation of synapse density for superficial (position 1) and middle layers (position 3) indistinguishable during adolescence (Fig. 7B,C) (P35 relative maturation PFC Position 1: 0.73 ± 0.14; Position 3: 0.76 ± 0.16; p = 0.4).

**Figure 7.**
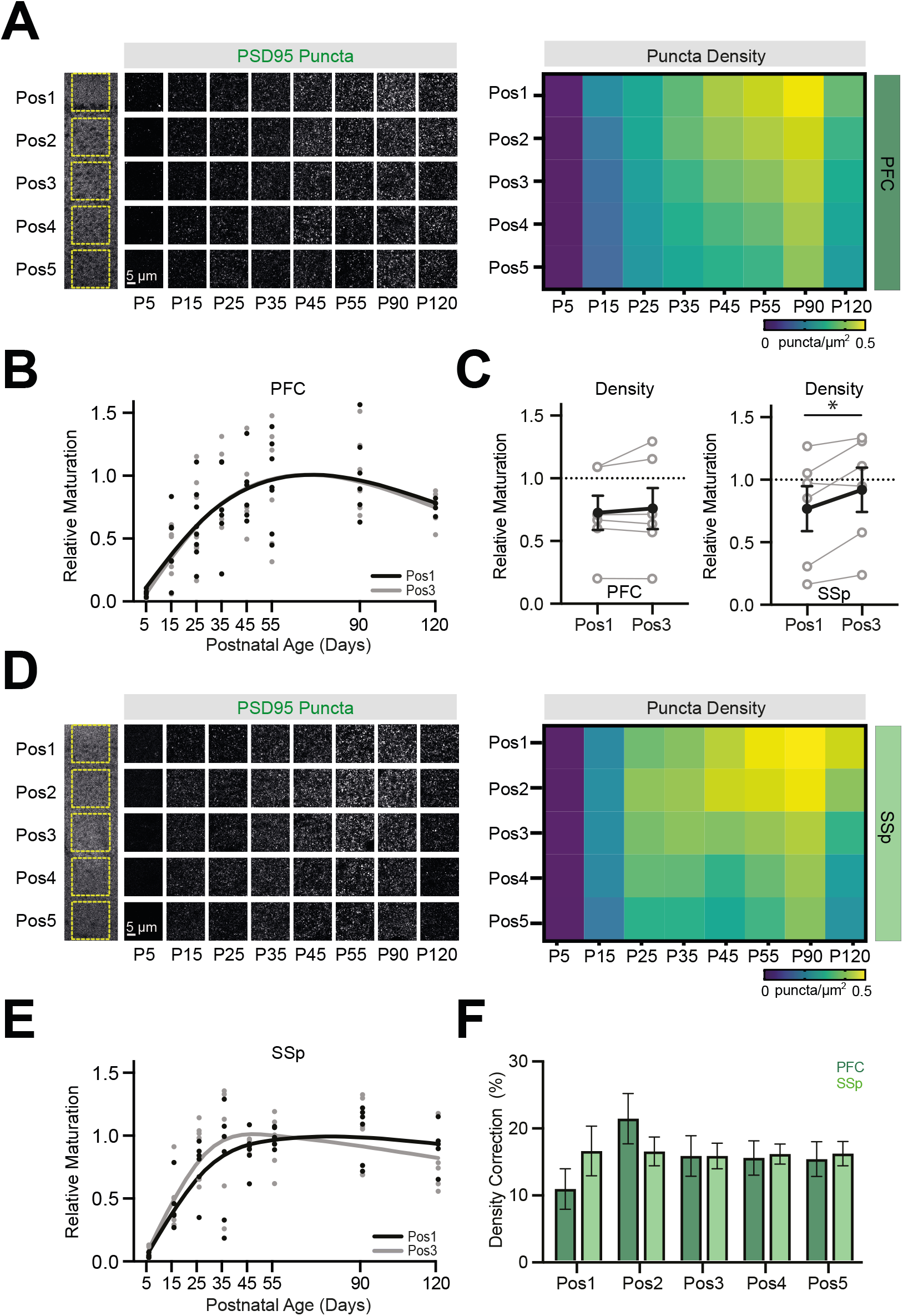
A) (*Left*) Representative high magnification images from the PFC showing PSD95 puncta labelling at positions 1-5 spanning the cortical depth. (*Right*) Heatmap showing average PSD95 puncta density at each PFC imaging position across development. B) Relative maturation of average puncta density in PFC at position 1 (Pos1) and position 3 (Pos3) normalized to adult levels (P90). Points represent the average puncta density for individual mice with smoothing spline fit to the individual data points. C) (*Left*) Relative PFC PSD95 puncta density in position 1 and position 3 at postnatal day (P)35 showing pairwise, within-slice comparisons. (*Right*) Relative SSp PSD95 puncta density in position 1 and position 3 at postnatal day (P)35 showing pairwise, within-slice comparisons. D) As **A** but for SSp. E) As **B** but for SSp. F) Percentage correction in PSD95 puncta density due to nuclear area across the different imaging positions in PFC and SSp. Number of mice per age group: PFC, P5 = 6, P15 = 6, P25 = 8, P35 = 6, P45 = 6, P55 = 7, P90 = 6, P120 = 5; SSp, P5 = 6, P15 = 6, P25 = 6, P35 = 6, P45 = 6, P55 = 6, P90 = 6, P120 = 5. Values are shown as mean average ± SEM, with exception of data in B & E which show smoothing spline fit to the individual data points. * = p < 0.05.

In SSp we found that puncta density was highest in middle layers (position 3) until P35. Subsequent late maturation of superficial layers meant that by adulthood there was greater synapse density in positions 1 and 2 (Fig. 7D). Consistent with the thalamus first development observed in SSp, the relative maturation of puncta density in layer 4 (position 3), matured earlier than superficial layers (position 1) (Fig. 7C-F) (P35 relative maturation SSp Position 1: 0.77 ± 0.18; Position 3: 0.92 ± 0.18; p = 0.03). When correcting for differences in neuronal density we found largely similar results (Fig. 7F & S8). For both regions we performed similar analysis for puncta size and mean intensity (Fig. S9). In SSp puncta size mirrored puncta density, showing a significant bias towards earlier maturation in position 3, which overlaps with L4 (P35 relative maturation SSp Position 1: 0.95 ± 0.05; Position 3: 0.99 ± 0.04; p = 0.006), while mean intensity developed at a similar rate across layers (P35 relative maturation SSp Position 1: 0.8 ± 0.16; Position 3: 0.87 ± 0.17; p = 0.15). By contrast, in PFC both synapse size (P35 relative maturation PFC Position 1: 0.93 ± 0.03; Position 3: 0.95 ± 0.03; p = 0.002) and intensity (P35 relative maturation PFC Position 1: 0.78 ± 0.11; Position 3: 0.83 ± 0.12; p = 0.02) showed a small yet significant bias towards maturing earlier in deeper layers.

In summary, these findings corroborate our earlier widefield imaging experiments, highlighting key differences in the way that PFC and SSp develop. In the adult, individual layers display notable differences in relative puncta density, with both regions showing enrichment of puncta in supragranular layers, mirroring previous results in adult mice (Zhu et al., 2018). However, individual layers largely develop in unison across PFC, while in SSp there is a sequential maturation with granular and infragranular layers developing earlier than supragranular layers. Changes in PSD95 levels are driven by an increasing number of synapses that contain more PSD95 at older ages.

### Modelling synapse dynamics in the developing cortex

In contrast to previous studies of dendritic spine density (Bourgeois et al., 1994; Pöpplau et al., 2024), we did not observe any adolescent pruning of PSD95 puncta across our analyses. We reasoned that this could be because not all spines, particularly immature ones, contain functional synapses or express PSD95 (Arellano et al., 2007; Knott et al., 2006; Zhu et al., 2018) and that the proportion of immature and mature spines is highly dynamic across development (Holtmaat et al., 2005). Consequently, our PSD95 data may better model the emergence of mature, stable synapses in the developing brain, rather than the total synapse population. To test this hypothesis, we built a computational model of synapse dynamics which incorporated previous *in vitro* and *in vivo* studies (see methods). At each stage of development individual synapses occupied either a reserve pool of potential synapses, a population of immature synapses, or a population of mature synapses (Fig. 8A). We first modelled synapse dynamics using constant rates fixed across development to govern the transition between these populations. We tested two versions of the model: a random walks model and a differential equation model, which yielded similar results (Fig. 8C & S10). Interestingly, we found that, although we were able to generate developmental trajectories for mature synapses that mirrored our PSD95 data using these models (Fig. 8C), pruning of the total synapse population was mathematically impossible if rates were fixed across development and the total synapse count begins at zero (Fig. 8C & proof in appendix 1). Next, we applied dynamic rates across development (Fig. 8C & S10), consistent with *in vivo* data on spine dynamics in the postnatal neocortex that show how the rate of creation and elimination decreases with age (Holtmaat et al., 2005). Under these conditions the three synapse populations displayed distinct trajectories during development. The immature population peaked early (<P25) before decaying to reach a stable level in adulthood. The mature population mirrored our observed developmental profile of PSD95, displaying a slow monotonic increase before stabilising towards the end of adolescence. Finally, the total synapse population peaked prior to adolescence, before undergoing a developmental reduction (Fig. 8C).

**Figure 8.**
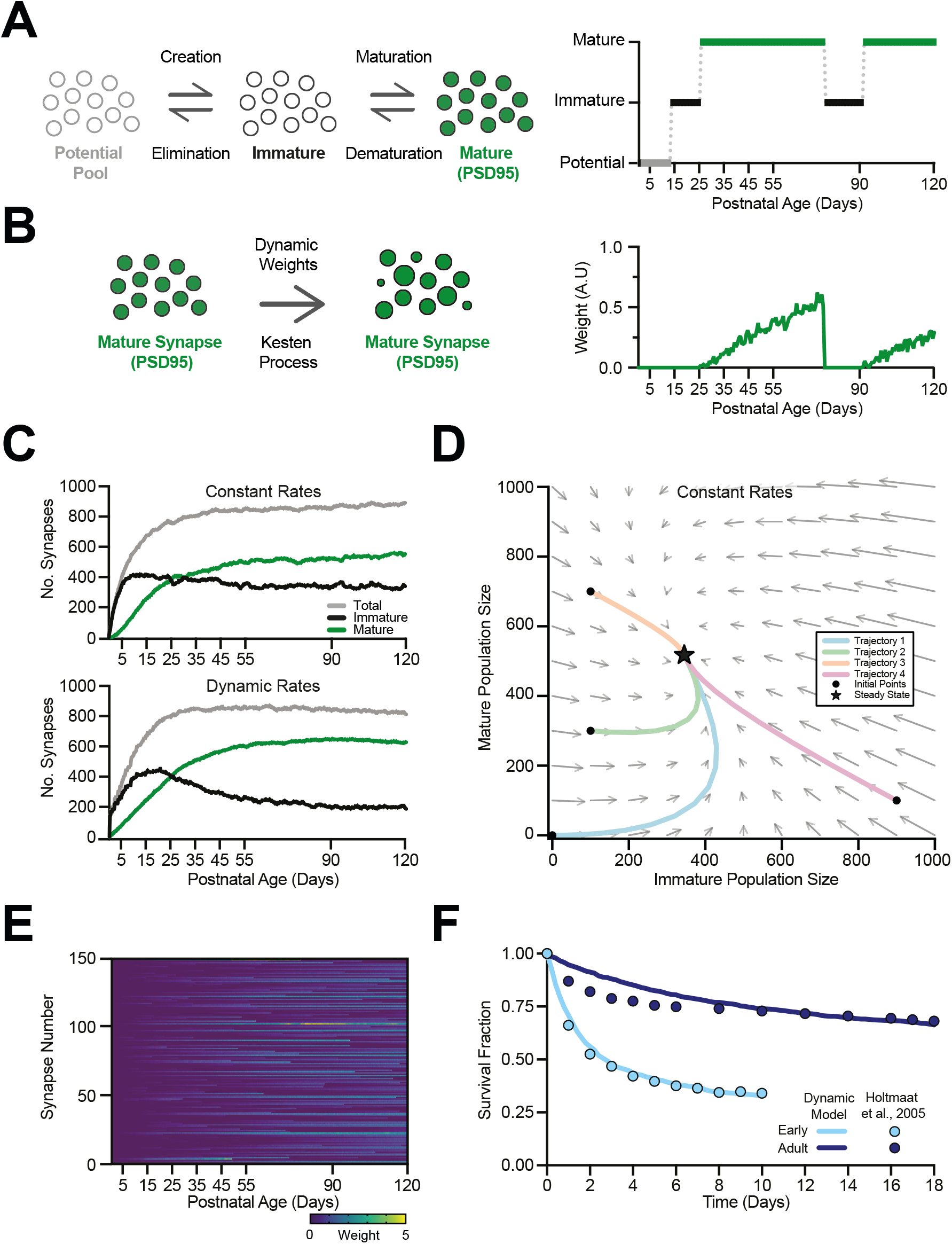
A) (*Left*) Model schematic showing how synapses move between potential, immature and mature populations. (*Right*) Example time series of a single synapse’s state during a simulation of 120 postnatal days, showing stochastic transitions between populations across development. B) (*Left*) Schematic showing how the mature synapses’ weights are dynamically modulated during development based on a Kesten Process (see methods). (*Right*) Example synapse weight showing stochastic changes across development, same simulation as panel A. C) (*Top*) Random walks model with constant rates governing the transition between synapse populations across development. Changes in immature, mature and total synapse number are shown. (*Bottom*) Similar but for model with dynamic weights across development (see **Fig. S10B**). D) Phase plot calculated from the differential equation model with constant rates, showing convergence of developmental trajectories in synapse number towards a fixed final state, despite different initial starting conditions. E) Heatmap showing lifetime synapse weight dynamics across development for a representative simulation of a population of 150 individual synapses, showing both persistent and transient synapses across development. F) Survival dynamics of synapses relative to two different reference ages: early (P16) and adult (P70). The plot compares results from the dynamic rates model (solid curves) and experimental data from (Holtmaat et al., 2005) (filled circles).

When synapses reached the mature population, their weight was also dynamically adjusted (Fig. 8B), with larger synapses resistant to de-maturation (Knott et al., 2006; Taft and Turrigiano, 2014) (Fig. S10). We modelled synapse lifetimes across development, observing both short-lived synapse and stable synapses that persisted across development (Fig. 8E). By computing the survival fraction for synapses at early (P16) and adult (P70) developmental ages we were able to show that immature synapses were less stable than adult synapses (Fig. 8F), mirroring previous *in vivo* work (Holtmaat et al., 2005). These results highlight potential mechanisms that govern synapse dynamics in the developing cortex. To determine how individual model parameters influence the properties of this system we performed sensitivity analysis. This revealed a stark separation in the developmental roles of each part of the model. The rates of synapse creation and elimination govern the developmental age at which the peak of the different synapse populations occurs, while the maturation and de-maturation rates govern the final number of mature and immature synapses (Fig. S10). Overall, these modelling results reconcile prior findings with our new data, showing developmental pruning of total dendritic spine number and a monotonic increase in mature (putative PSD95+) synapses.

## DISCUSSION

This study maps the development of glutamatergic synapses across layers and regions of the mouse neocortical hierarchy by quantifying changes in fluorescently-labelled MAGUK synaptic scaffold proteins PSD95 and SAP102. We find key differences in the way that PSD95 expression develops in primary sensory and high-order prefrontal areas, with sensory cortex development occurring earlier and in a layer-specific manner. By contrast, prefrontal development occurs later and is largely synchronous across layers. This relationship is restricted to PSD95, with SAP102 displaying earlier expression and few layer-dependent effects across the cortex. Adolescence is known to be important for PFC maturation, and we observe pronounced changes in PFC synapses across all layers during this time. However, in SSp we also observe significant maturation in superficial synapses, particularly those in L1. Cortical L1 is a key locus for the integration of higher-order, top-down inputs (Huang et al., 2024; Schuman et al., 2021), such that our findings are consistent with a model whereby feedforward pathways develop early, followed by the gradual maturation of higher-order areas and their feedback projections, culminating in the final stage of inside-out development. Importantly, we find that this change is observed across much of cortex, suggesting adolescence represents an important period for the development of top-down inputs.

Adolescence is a key time point for the prefrontal cortex, with changes observed at the level of connectivity, receptor expression, and intrinsic physiology resulting in refinements to network dynamics and cognition (Anastasiades et al., 2022; Delevich et al., 2019; Larsen and Luna, 2018; Pöpplau et al., 2024; Zhu et al., 2024). Our findings are consistent with the idea that glutamatergic synapses in PFC increase the relative occupancy and amount of PSD95 synaptic protein during adolescence. This likely helps stabilize synaptic connections and promote their lifetime (Bulovaite et al., 2022; Delevich et al., 2019; Ehrlich et al., 2007) while also providing a structural scaffold for key components of the synaptic signalling machinery (Lohmann and Kessels, 2014; Sheng and Kim, 2011). Our findings support the idea that this process occurs at the same time across layers. However, it is important to note that each layer contains many different cell types, and although certain inputs are biased towards a given layer, they may target specific neurons across distinct parts of their dendritic arbour (Anastasiades and Carter, 2021). This is also true for the adolescent maturation of synapses we observe in L1, with previous studies highlighting cell-type-specific connectivity (Anastasiades et al., 2021), developmental dynamics and plasticity in this layer (Ibrahim et al., 2021; Klappenbach et al., 2025). Our findings are therefore not inconsistent with input-or cell-type-specific effects, although it appears that, at least at the level of layers, such a process is less evident in PFC than in sensory and motor areas.

The protracted maturation of top-down feedback to L1 is consistent with a model whereby the cortex gradually transitions from an early feedforward dominated network to one where feedback projections take on greater prominence. This phenomenon has recently been reported in human *in vivo* studies, with top-down activity also elevated during cognitively demanding tasks (Pines et al., 2023). The increased prominence of feedback connectivity has been proposed to mark the developmental inflection point at which the brain transitions from childhood to adolescence (Dong et al., 2021) and occurs contemporaneously with the maturation of both the cortical neurons providing feedback (Nabel et al., 2020) and the higher-order thalamo-cortical system (Sydnor et al., 2025). Top-down connections have increasingly been incorporated into brain-inspired machine learning models that use feedback as a teaching signal that is propagated from higher to lower levels of the network (Sacramento et al., 2018; Tugsbayar et al., 2025). In such models it is often advantageous to use a dynamic, or adaptive, learning rate to help optimize the network. It is interesting to speculate that the brain may also employ a similar strategy, relying more heavily on feedforward information during early development but employing increasingly sophisticated internal predictions and outcome monitoring performed by higher-order cortices to guide learning as development ensues (Buzzell et al., 2017; Hartley and Somerville, 2015). Consistent with this idea, L1 is a key site of integration for top-down multi-modal signals (Huang et al., 2024; Schuman et al., 2021) that help regulate learning, sensory encoding, perceptual decision making, and flexible behaviour (Banerjee et al., 2020; Doron et al., 2020; Furutachi et al., 2024; Letzkus et al., 2011; Mo et al., 2024; Williams and Holtmaat, 2019). Importantly, weak top-down signalling is observed in schizophrenia, a neurodevelopmental disorder that emerges during adolescence and whose symptomatology includes impairments in learning and decision making (Lyu et al., 2025). The maturation of L1 may therefore represent a key locus for the emergence of flexible cognition and, consequentially, a key site of vulnerability in adolescent mental health disorders.

The distinct developmental timings we observe in SSp versus PFC are consistent with the hierarchical maturation of individual brain areas, with early sensory regions maturing before association areas (Gogtay et al., 2004; Pinto et al., 2013; Reh et al., 2020). However, there is evidence from primates that synaptogenesis occurs concurrently throughout much of cortex, with synapse density peaking prior to puberty, before undergoing a period of pruning during adolescence (Bourgeois et al., 1994; Zecevic and Rakic, 1991) and plasticity in SSp is readily observed in non-granular layers during adolescence, but less so thereafter (Fox, 2002). Although our data support the hierarchical development of cortex, we also see adolescent changes in PSD95 levels across much of cortex. Contrary to the decrease in synapse number expected in response to pruning, we instead observe a greater number of PSD95 positive puncta, consistent with previous studies using this mouse line (Cizeron et al., 2020). A similar disconnect between synapse number and synaptic protein expression has been reported for frontal regions across species, with synapse elimination observed in adolescent humans (Huttenlocher and Dabholkar, 1997), primates (Bourgeois et al., 1994) and rodents (Pöpplau et al., 2024) despite bulk proteome studies displaying a steady increase in the levels of PSD95 and other synaptic markers during the same period (Glantz et al., 2007; Pinto et al., 2013; Webster et al., 2011). This discrepancy has led to a debate in the literature with regards synaptic pruning in the neocortex (Webster et al., 2011). Our computational modelling provides a way to reconcile these data whereby nascent, immature synapses, which typically lack or express low levels of PSD95, are produced at high rates early in development, peaking just prior to adolescence. The dynamic developmental shifts in the rates of spine formation and elimination cause subsequent reduction in spine number (Holtmaat et al., 2005), while at the same time mature synapses incorporate PSD95 clusters of increasing size. This is consistent with cortex-wide data showing how in the nascent cortex synapses are predominantly SAP102 positive, while over the course of the first postnatal weeks there is a gradual increase in the proportion of PSD95 or SAP102/PSD95 positive synaptic puncta. This occurs concomitantly with the maturation of synaptic molecular diversity, which also begins to stabilize towards the end of adolescence (Cizeron et al., 2020).

Integrating these data, it is possible to explain the mechanisms through which changes in glutamatergic synapses emerge across the rodent neocortex. PSD95 is often associated with glutamatergic spines, however the two are not ubiquitous, spines can form even after complete loss of PSD95 (Yusifov et al., 2021), PSD95 may be present on aspiny GABAergic interneurons (Kawabata et al., 2012), and cortical synapses are highly diverse and can be enriched for distinct MAGUK proteins (Zhu et al., 2018). This process is also highly developmentally regulated. The early neocortex is dominated by immature “silent” synapses which primarily signal via NR2B subunits of the NMDA-R (Hanse et al., 2013). The NR2B subunit couples to SAP102 to promote synapse formation (Chen et al., 2011). This is consistent with higher levels of SAP102 and very low levels of PSD95 in both PFC and SSp prior to P15 and observations that the early synaptic maturation of these brain regions is similar (Myme et al., 2003). At postnatal day 12 the brain undergoes a period of sensory awakening which coincides with eye opening, active whisking and the unblocking of the auditory canal (Wu et al., 2024). This drives the rapid localization of PSD95 to the synapse (Yoshii et al., 2003), which we observe to increase at P15. This shift facilitates synapse maturation via activity dependent AMPA-R insertion and the transition to NR2A mediated NMDA-R signalling (Elias et al., 2008; Hanse et al., 2013). Over time this causes an abundance of nascent synapses, which peak in the days following (Holtmaat et al., 2005). As development ensues synapses continue to incorporate PSD95 while a subset of synapses increase their size and stability. Synapses that fail to undergo stabilization or downregulate PSD95 expression are lost (Holtmaat et al., 2005; Knott et al., 2006). This process likely contributes to the termination of developmental plasticity (Huang et al., 2015).

Our modelling of spine dynamics replicates the adolescent changes in PSD95 puncta observed in both PFC and L1 of sensory cortices, as well as *in vivo* observations (Holtmaat et al., 2005). Our model suggests that maturing PSD95 puncta may act as a “developmental clock” tracking the relative maturity and stability of synapses in different brain regions. Although various biological mechanisms can control the rates of synapse elimination, maturation and de-maturation (Yoshihara et al., 2009), it remains unclear exactly how these rates may vary across brain regions and synapse types; differences in neural activity (Zuo et al., 2005), synaptic/structural proteins (Agoglia et al., 2017; Zhu et al., 2018), cytoskeletal dynamics (Honkura et al., 2008), sex hormone receptors (Delevich et al., 2019; McEwen and Milner, 2017), or microglial pruning (Pöpplau et al., 2024; Schalbetter et al., 2022) may play a role by fine tuning the rates at which synapses transition between different states to yield region and layer-specific trajectories. It will be important to determine the mechanisms through which synapse maturation is orchestrated given significant differences exists, even within a single brain region.

In summary, we observe adolescence to be a key period for the maturation of synapses associated with higher-order circuits in the neocortex, which includes all layers of association regions and L1 of lower-order cortices, including primary sensory regions. These changes occur due to increases in the number of synapses incorporating PSD95 molecules as well as likely increases in synapse strength and stability. The patterns of synaptic maturation we observe are consistent with enhanced cognitive capabilities during adolescence, including improved top-down control of behaviour (Dong et al., 2021; Hartley and Somerville, 2015; Pines et al., 2023). Our current study only focuses on individual synapses, without identifying their source or postsynaptic target. In the future, understanding the mechanisms and specificity that governs how these circuits mature will advance our understanding of healthy aging and may also be of clinical significance given adolescence is a key period for the emergence of various mental health conditions (Blakemore, 2019; Solmi et al., 2021).

## Acknowledgements

We would like to thank members of the Ashby, Anastasiades, and Mellor labs for helpful discussions. Imaging was conducted in the Wolfson Bioimaging Facility at the University of Bristol. We would particularly like to thank Dr Stephen Cross who assisted with image analysis pipelines and Katy Jepson who provided support for slide scanner imaging.

## Contributions

Data collection: LD, JM, RR, SA, GM-S, DF. Data analysis: LD, JM, RR, SA, GM-S, PGA. Manuscript writing: PGA, LD, JM, RR. Manuscript editing: All authors. Supervision: PGA, MA, CO’D. Funding Acquisition: PGA, MA, CO’D, SGNG.

## Supplemental Figure Legends

**<u>Figure S1</u>**

Relates to figure 1.

A) Relative maturation of PSD95 and SAP102 at P5 compared to adult (P90) levels. Showing pairwise comparison in the PFC (*left*) and SSp (*right*).

B) SAP102/PSD95 ratio for PFC and SSp at different developmental ages highlighting how changes in the relative expression of the two proteins occur during development.

C) Inter-regional correlational in PSD95 levels between PFC and SSp in individual mice across different developmental ages.

D) Inter-regional correlational in SAP102 levels between PFC and SSp in individual mice across different developmental ages.

Values are shown as mean average ± SEM. Data in B show smoothing spline fit to the individual data points. * = p < 0.05.

**<u>Figure S2</u>**

Relates to figures 2 & 3.

A) (*Left*) Representative images of VGlut2 staining across PFC layers at P5, P15 and P55. (*Right*) Average VGlut2 fluorescent profiles across all mice imaged at P5, P15 and P55 showing shift in distributions across development. Dashed lines indicate individual layer boundaries.

B) Quantification of the cortical depth of the second peak in VGlut2 labelling, corresponding to the location of L3 in the PFC (Collins et al., 2018). Data is shown at different postnatal ages spanning cortical development.

C&D) Similar to **A&B** but for SSp. Data in D calculates the cortical depth corresponding to the start of the L4 barrels.

Values are shown as mean average ± SEM, with exception of plots in A & C where error bars have been omitted for clarity. * = p < 0.05.

**<u>Figure S3</u>**

Relates to figures 2 & 3.

A) Representative images of DAPI staining across PFC layers during postnatal development. Dashed lines indicate individual layer boundaries.

B) Representative high magnification images of DAPI labelling at different positions spanning the depth of the PFC at P5 and P15. Note the reduction in cell density that occurs between the two ages.

C&D) As **A&B** but for SSp.

E) Heatmap showing average cell density at different positions spanning the cortical depth in PFC (left) and SSp (right) during postnatal development.

Values are shown as mean average.

**<u>Figure S4</u>**

Relates to figures 4.

A) Absolute adolescent change in SAP102 fluorescence levels calculated by subtracting the P55 values by the P25 values. Data points show average change for individual layers across cortical regions.

B) Absolute adolescent PSD95 change for individual cortical layers. Each data point represents the average change observed in a single brain region.

C) Ratio of adolescent change in PSD95 fluorescence levels calculated by first dividing the value for each layer (LX) by the corresponding value in L6 of an individual brain region and age to generate a within mouse ratio at P25 and P55. These ratios are then divided (P55/P25) to give a developmental ratio. Data points show average change for L1, L2/3 and L5 across cortical regions.

D) Correlation of the LX/L6 PSD95 ratio change with the hierarchy position of each cortical area. Data is shown for L1, L2/3 and L5.

Values are shown as mean average ± SEM, with exception of plots in A,C & D where error bars have been omitted for clarity. * = p < 0.05.

**<u>Figure S5</u>**

Relates to figures 5.

A) Principal component plot highlighting two clusters based on raw PSD95 expression levels at P55. Cluster 1 (blue) contains primarily sensory-motor regions and cluster 2 (red) contains mostly higher-order association cortices.

B) Heatmap showing the absolute PSD95 expression by layer and regions at P55. X indicates agranular cortex lacking L4.

C) Comparison of the relative adolescent change in L5 (*left*) and L6 (*right*) PSD95 signal between cluster 1 and cluster 2.

E) Correlation of the LX/L6 PSD95 ratio change with the hierarchy position of individual visual cortical area.

E) Absolute PSD95 levels in L1 and L2/3 for cluster 1 and cluster 2 at P25.

F) Absolute PSD95 levels in L1 and L2/3 for cluster 1 and cluster 2 at P55.

Values are shown as mean + SEM. SEM are omitted in B & D for clarity. * = p < 0.05.

**<u>Figure S6</u>**

Relates to figure 6.

A) (Top) Representative high magnification image showing PSD95 and DAPI labelling. (Bottom) Cropped region of interest corresponding to the region of interest highlighted in the top image.

B) Examples of puncta detection performed on the highlighted region of interest in A (bottom). Differences values for the puncta radius (R) and threshold (T) shift the number of puncta detected. Common puncta across all 3 settings shown in black, false positives in the high detection setting in red (top) and false negatives in the low detection setting in blue (bottom).

C) Heat map showing PSD95 puncta density from a subset of images sampled at different developmental ages. Red, yellow, and blue boxes highlight the settings used in **B** for high, optimal and low detection, respectively. Note that although the absolute number of puncta detected varies across the different detection settings the developmental increase in puncta density occurs across regardless of the detection settings.

D) Number of PSD95 puncta detected at P5 using three different laser intensities. 1x represents the value used for the main study which ensured reliable puncta detection at adult ages without oversaturated images. Increasing the laser intensity to 2x or 2.5x yielded a slight increase in synapse number at P5 but remained much less than the number of synapses observed at P15 using 1x intensity.

E) Relative PFC (*left*) and SSp (*right*) maturation calculated based on widefield fluorescence values (10x) and using a model which multiplies the puncta density x intensity x size for each mouse at each age (Model).

Values are shown as mean average ± SEM. Data in E show smoothing spline fit to the individual data points.

**<u>Figure S7</u>**

Relates to figure 7.

A) Schematic showing how each high magnification imaging position overlaps with the individual layers of the PFC at P5 (left) and ages from P15 onwards (right).

B) Similar to **A** but for SSp.

**<u>Figure S8</u>**

Relates to figures 7.

A) Heatmap showing average nuclear area at different positions spanning the cortical depth in PFC (left) and SSp (right) during postnatal development.

B) Heatmap showing average PSD95 puncta density after correcting for nuclear area at different positions spanning the cortical depth in PFC (left) and SSp (right) during postnatal development.

Values are shown as mean average.

**<u>Figure S9</u>**

Relates to figures 7.

A) (*Top*) Heatmap showing average PSD95 puncta size at each PFC imaging position across development. (*Bottom*) Relative maturation of average puncta size in PFC at position 1 (Pos1) and position 3 (Pos3) normalized to adult levels (P90). Points represent the average puncta size across the cortical depth for individual mice with smoothing spline fit to the individual data points.

B) Similar to **A** but for PFC puncta mean intensity. C&D) Similar to **A&B** but for SSp.

Values are mean average. Data in **A-D** show smoothing spline fit to the individual data points.

**<u>Figure S10</u>**

Relates to figure 8.

A) (*Left, right*) Graphs showing overlay of random walks and differential equations model with constant rates (left) or dynamic rates (right) governing the transition between synapse populations across development. Colours indicate immature (black), mature (green) and total (grey) synapse population counts. In all cases the random walk and differential equation models give very similar results.

B) (*Top*) Creation (red) and elimination (blue) rates at different developmental ages in the dynamic rates model. (*Bottom*) De-maturation rate (purple) as a function of synapse weight. Bigger synapses are more stable and more resistant to de-maturation.

C) Sensitivity analysis of how model parameters governing the transition rates in the dynamic rates model (columns of matrix) influence the eventual number of mature or immature synapses at P120 (top two rows of matrix), as well as the developmental age that the peak density of the three different synapse types is reached (bottom three rows of matrix). *A*_1_, *A*_2_ are P0 baseline offsets, and *τ*_1_, *τ*_2_ are decay time constants dictating decline of creation and elimination rates with developmental age, respectively. *m* is maturation rate, *A*_3_ and *γ* are zero-weight amplitude and weight-decay constant dicatating weight-dependence of of de-maturation rates, respectively.

