## Supplemental Appendix for "Region- and layer-specific glutamatergic synapse development in the nascent cortical hierarchy"

### 1 Appendix

#### 1.1 Analytical solution to differential equations with constant rates

The differential equations

$$\frac{dN_I}{dt} = cN_P - (e+m)N_I + dN_M$$
$$\frac{dN_M}{dt} = mN_I - dN_M$$
$$\frac{dN_P}{dt} = eN_I - cN_P$$

can be rewritten as

$$\frac{dN_I}{dt} = cT - (e+m+c)N_I + (d-c)N_M$$
$$\frac{dN_M}{dt} = mN_I - dN_M$$

where $T$ is the *Total* possible synapses, $T = N_P + N_I + N_M$.

This system of ordinary differential equations can be written as

$$\vec{s'}(t) = \frac{d}{dt}\begin{bmatrix} N_I \\ N_M \end{bmatrix} = \begin{bmatrix} -(e+m+c) & (d-c) \\ m & -d \end{bmatrix}\begin{bmatrix} N_I \\ N_M \end{bmatrix} + \begin{bmatrix} cT \\ 0 \end{bmatrix}$$

An analytical solution of the form $\vec{s}(t) = \vec{s_h}(t) + \vec{s_p}(t)$ can be found, where $\vec{s_h}$ and $\vec{s_p}$ denote the homogeneous and particular solutions, respectively. The solution to the homogeneous case is in the form

$$\vec{s_h}(t) = c_1\vec{v_1}e^{\lambda_1 t} + c_2\vec{v_2}e^{\lambda_2 t}$$

where $\lambda_1, \lambda_2$ are the eigenvalues of the system matrix, $\vec{v_1}, \vec{v_2}$ the eigenvectors, and $c_1, c_2$ constants. The eigenvalues are

$$\lambda_i = \frac{-(e+m+c+d) \pm \sqrt{(e+m+c+d)^2 - 4(ed+dc+mc)}}{2}$$

and the eigenvectors

$$\vec{v_i} = \begin{bmatrix} 1 \\ \frac{m}{d+\lambda_i} \end{bmatrix}$$

The constants $c_1, c_2$ can then be determined from the initial values, i.e. $N_I(0) = 0, N_M(0) = 0$. We get that

$$c_1 = \frac{-\lambda_2 cT(d+\lambda_1)}{(\lambda_1-\lambda_2)(ed+cd+cm)}, \quad c_2 = \frac{-\lambda_1 cT(d+\lambda_2)}{(\lambda_2-\lambda_1)(ed+cd+cm)}$$

For the particular solution (the inhomogeneous case), we use the method of undetermined coefficients. Given that the inhomogeneous term is $\begin{bmatrix} cT \\ 0 \end{bmatrix}$, we let $\vec{s_p}(t) = \vec{k} = \begin{bmatrix} k_1 \\ k_2 \end{bmatrix}$, which gives $\vec{s_p}' = \begin{bmatrix} 0 \\ 0 \end{bmatrix}$ Substituting this into our differential equation for $\vec{s'}$ gives

$$\begin{bmatrix} 0 \\ 0 \end{bmatrix} = \begin{bmatrix} -(e+m+c) & (d-c) \\ m & -d \end{bmatrix}\begin{bmatrix} k_1 \\ k_2 \end{bmatrix} + \begin{bmatrix} cT \\ 0 \end{bmatrix}$$

This allows us to solve for $k_1, k_2$, yielding

$$k_1 = \frac{dcT}{ed+cd+cm}, \quad k_2 = \frac{mcT}{ed+cd+cm}$$

We now have a full expression for the solution:

$$\vec{s}(t) = c_1\vec{v_1}e^{\lambda_1 t} + c_2\vec{v_2}e^{\lambda_2 t} + \vec{k}$$

#### 1.2 Monotonically increasing combined population

We want to ascertain whether the overall, combined population $N_I + N_M$ necessarily increases monotonically (from the origin) with constant rates $c, e, m, d$, or if it is possible that an initial increase in $N_I + N_M$ can be followed by a decrease ("pruning back"). In order to analyse the combined population, we take the analytical solution $\vec{s}(t) = c_1\vec{v_1}e^{\lambda_1 t} + c_2\vec{v_2}e^{\lambda_2 t} + \vec{k}$, and sum the two rows of the system of equations, i.e.

$$\begin{aligned}\text{Combined population (C.P)} &= c_1v_{11}e^{\lambda_1 t} + c_2v_{21}e^{\lambda_2 t} + k_1 + c_1v_{12}e^{\lambda_1 t} + c_2v_{22}e^{\lambda_2 t} + k_2 \\ &= c_1e^{\lambda_1 t}(v_{11}+v_{12}) + c_2e^{\lambda_2 t}(v_{21}+v_{22}) + k_1 + k_2\end{aligned}$$

To test for non-monotonicity (an increase in $N_I + N_M$ followed by a decrease), we find the derivative of the expression:

$$\frac{d}{dt}\text{C.P.} = \lambda_1c_1e^{\lambda_1 t}(v_{11}+v_{12}) + \lambda_2c_2e^{\lambda_2 t}(v_{21}+v_{22})$$

In order for there to be a local maximum, there must be a $t$ such that the derivative is equal to 0. Setting it equal to 0 gives

$$\lambda_1c_1e^{\lambda_1 t}(v_{11}+v_{12}) = -\lambda_2c_2e^{\lambda_2 t}(v_{21}+v_{22})$$

$$\Rightarrow \frac{e^{\lambda_1 t}}{e^{\lambda_2 t}} = e^{(\lambda_1-\lambda_2)t} = \frac{-\lambda_2c_2(v_{21}+v_{22})}{\lambda_1c_1(v_{11}+v_{12})}$$

We solve for $t$:

$$(\lambda_1-\lambda_2)t = \ln(\kappa) \Rightarrow t = \frac{\ln(\kappa)}{\lambda_1-\lambda_2}$$

where

$$\kappa = \frac{-\lambda_2c_2(v_{21}+v_{22})}{\lambda_1c_1(v_{11}+v_{12})}$$

Therefore, $t$ exists when $\kappa > 0$. In order to disprove that the combined population increases, reaches a maximum, and then decreases, we need to show that $\kappa < 0$ always. Let us substitute in the expressions of $c_1, c_2$ presented above.

$$\Rightarrow \kappa = \frac{-\lambda_2(\frac{-\lambda_1cT(d+\lambda_2)}{(\lambda_2-\lambda_1)(ed+cd+cm)})(v_{21}+v_{22})}{\lambda_1(\frac{-\lambda_2cT(d+\lambda_1)}{(\lambda_1-\lambda_2)(ed+cd+cm)})(v_{11}+v_{12})} = \frac{-(d+\lambda_2)(\lambda_1-\lambda_2)(v_{21}+v_{22})}{(d+\lambda_1)(\lambda_2-\lambda_1)(v_{11}+v_{12})}$$

Now let us write in the terms for $v_{11}, v_{12}, v_{21}, v_{22}$:

$$\kappa = \frac{-(d+\lambda_2)(\lambda_1-\lambda_2)(1+\frac{m}{d+\lambda_2})}{(d+\lambda_1)(\lambda_2-\lambda_1)(1+\frac{m}{d+\lambda_1})} = \frac{-(\lambda_1-\lambda_2)(d+m+\lambda_2)}{(\lambda_2-\lambda_1)(d+m+\lambda_1)}$$

The part of the expression $\frac{(\lambda_1-\lambda_2)}{(\lambda_2-\lambda_1)}$ is $-1$, which means

$$\kappa = \frac{(d+m+\lambda_2)}{(d+m+\lambda_1)}$$

We now prove the eigenvalues of this system are real by showing that the discriminant $(e+m+c+d)^2 - 4(ed+dc+mc) \geq 0$. We claim this is greater than or equal to 0:

$$\text{Claim: } (e+m+c+d)^2 - 4(ed+dc+mc) \geq 0$$

First, we note that $(e+m+c+d)^2 - 4(ed+dc+mc)$ can be rewritten as

$$(d-c)^2 + (e+m)^2 - 2(d-c)(e-m).$$

This means that

$$(e+m+c+d)^2 - 4(ed+dc+mc) \geq 0 \Leftrightarrow (d-c)^2 + (e+m)^2 \geq 2(d-c)(e-m).$$

We note that if $d = c$, then it is obviously true. If $d > c$, then

$$(d-c)^2 + (e+m)^2 \geq 2(d-c)(e+m) \geq 2(d-c)(e-m).$$

The first inequality follows since $x^2+y^2 \geq 2xy$. The second inequality follows since $d-c > 0$ and $e+m > e-m$. From this it follows that

$$(d-c)^2 + (e+m)^2 \geq 2(d-c)(e-m), \text{ for } d \geq c.$$

To complete the proof, consider the case when $d < c$. Then

$$(d-c)^2 + (e+m)^2 \geq 2(c-d)(e+m) \geq 2(c-d)(m-e) = 2(d-c)(e-m).$$

The first inequality follows as when we re-arrange the expression and move $2(c-d)(e+m)$ to the left, we get $(d-c)^2 + (e+m)^2 - 2(c-d)(e+m)$, which can be rewritten as $(d-c)^2 + (e+m)^2 + 2(d-c)(e+m)$. This is a perfect square $(d-c+e+m)^2$, which is clearly non-negative, meaning that $(d-c)^2 + (e+m)^2 \geq 2(c-d)(e+m)$. The second inequality holds since $e+m \geq m-e$.

Thus we have proved that $(d-c)^2 + (e+m)^2 \geq 2(d-c)(e-m)$, and from this it follows that $(e+m+c+d)^2 - 4(ed+dc+mc) \geq 0$, which is the proof of our claim.

Substituting in the formulae we have for the eigenvalues into $\kappa$ yields

$$\kappa = \frac{(d+m+\frac{-(e+m+c+d)-\sqrt{(e+m+c+d)^2-4(ed+dc+mc)}}{2})}{(d+m+\frac{-(e+m+c+d)+\sqrt{(e+m+c+d)^2-4(ed+dc+mc)}}{2})}$$

$$= \frac{d+m-e-c-\sqrt{(e+m+c+d)^2-4(ed+dc+mc)}}{d+m-e-c+\sqrt{(e+m+c+d)^2-4(ed+dc+mc)}}$$

Multiplying top and bottom by the denominator's conjugate yields:

$$\kappa = \frac{\left(d+m-e-c-\sqrt{(e+m+c+d)^2-4(ed+dc+mc)}\right)^2}{(d+m-e-c)^2-(e+m+c+d)^2+4(ed+cd+cm)}$$

The numerator is always positive, therefore we need only check the denominator. Simplifying it gives $-4em$, and since $e, m$ are positive, $-4em$ is negative, making $\kappa < 0$. This concludes the proof. Therefore, with constant rates $c, e, m, d$ the combined population $N_I + N_M$ (with initial values $N_I(0) = N_M(0) = 0$) increases monotonically.
